# Systematic in vivo interrogation identifies novel enhancers and silencers associated to Atrial Fibrillation

**DOI:** 10.1101/2021.07.14.452222

**Authors:** Jesus Victorino, Isabel Rollan, Raquel Rouco, Javier Adan, Miguel Manzanares

## Abstract

Cis-regulatory elements control gene expression in time and space and their disruption can lead to pathologies. Reporter assays allow the functional validation of enhancers and other regulatory elements, and such assays by means of the generation of transgenic mice provide a powerful tool to study gene regulation in development and disease. However, these experiments are time-consuming and, thus, their performance is very limited. Here, we increase the throughput of in vivo mouse reporter assays by using a piggyBac transposon-based system, and use it to decode the regulatory landscape of atrial fibrillation, a prevalent cardiac arrhythmia. We systematically interrogated ten human loci associated to atrial fibrillation in the search for regulatory elements. We found five new cardiac-specific enhancers and implicated novel genes in arrhythmia through genome editing and three-dimensional chromatin analysis by 4C-seq. Of note, functional dissection of the 7q31 locus identified a bivalent regulatory element in the second intron of the *CAV1* gene differentially acting upon four genes. Our system also detected negative regulatory elements thanks to which we identified a ubiquitous silencer in the 16q22 locus that regulates *ZFHX3* and can outcompete heart enhancers. Our study characterizes the function of new genetic elements that might be of relevance for the better understanding of gene regulation in cardiac arrhythmias. Thus, we have .established a new framework for the efficient dissection of the genetic contribution to common human diseases.

## Introduction

The non-coding regulatory genome controls gene expression during development and homeostasis mainly through cis-regulatory elements (CREs), among which are positive elements such as enhancers (1). However, whereas hundreds of thousands of enhancers are predicted to exist in the human genome, only a small fraction have been functionally validated (2, 3).

Enhancer identification or validation usually comes from enhancer-reporter assays (ERAs) that allow testing changes in transcription driven by genomic sequences. ERAs are based on the use of a synthetic construct where the expression of a reporter gene is controlled by a minimal promoter with low levels of basal expression (4) that is delivered to living cells. If a genomic region with enhancer activity is inserted in the construct, there is a boost in transcription. However, while ERAs can be performed in tissue culture, the complexity of the regulatory genome and tissue-specificity of enhancers makes the selection of the right cell type and condition difficult (4). For instance, enhancers can be specifically active in a particular tissue during a precise developmental time point. Hence, testing a candidate enhancer in a single cell type shows a snapshot with reduced information that might lead to miss its potential activity.

On the other hand, the generation of transgenic animals harboring the ERA construct provides a powerful tool to study enhancer biology in all tissues at different developmental time points (5). Zygote microinjection with linear ERA constructs has been the classical way of assaying enhancers in mouse embryos (6–8). However, this procedure is time-consuming and has low efficiency, which limits the throughput of *in vivo* enhancer validation. Conversely, due to the success of comparative genomics and whole-genome epigenetic profiling, the list of putative enhancers to be tested has grown exponentially (9).

Genome-wide association studies (GWAS) have linked thousands of single-nucleotide polymorphisms (SNPs) in the non-coding genome to common diseases (10–12). Other forms of genetic variation such as copy number variants (CNV) or small insertion and deletions (indels) have also been linked to disease (13, 14). Although the most extended hypothesis is that many of these variants might be located at CREs such as enhancers, silencers or insulators, only a small number of these associations have been explored and experimentally tested (15, 16).

Atrial Fibrillation (AF) is the most common cardiac arrhythmia in humans (17). AF is now considered a polygenic condition (18, 19) and GWAS have identified over a hundred associations in AF patients as compared to healthy individuals (20–27). The first three loci and genes to be associated to AF through GWAS were *PITX2* at 4q25 (20), *ZFHX3* at 16q22 (21) and *KCNN3* at 1q21 (22), and they remain as the most significant associations found in all subsequent studies (23–27). However, a limitation of GWAS is that they do not have enough resolution to identify causal SNPs (28). Instead, the linkage disequilibrium (LD) block of the associated polymorphism is marked as a disease-risk locus. Since LD blocks span from a few to hundreds of kilobases (29–32), deciphering the CREs behind these associations has remained a challenge. As a result, the mechanism underlying the genetic component for an increased risk of AF remains elusive even for the first three loci identified more than a decade ago.

Here we have taken advantage of the piggyBac (PB) transposon system (33) to increase the throughput of classical transgenic-based ERA. Zygote microinjection of the PB-ERA system increased the efficiency of transgenesis to >60% and reproduced the expression patterns of previously characterized bona-fide enhancers. In order to prove PB-ERA as a useful tool to help dissect the functional genomics underlying common diseases, we focused on non-coding regions associated to AF. We interrogated ten human loci containing a total of 14 SNPs and 1 CNV and assayed their regulatory potential in mouse embryos. PB-ERA identified five cardiac-specific enhancers regulating *KCNIP1*, *C9orf3/AOPEP*, *SYNE2* and *CAV1*. We identified CRE targets and implicated new genes in AF through CRISPR/Cas9 genome editing and three-dimensional chromatin analysis by 4C-seq of human AF loci orthologous in mouse. In addition, we demonstrated the improved capabilities of the PB-ERA system to interrogate the regulatory genome with the identification of a ubiquitous silencer in the 16q22 locus controlling the expression of the transcription factor *ZFHX3*. Our study shows the value of in vivo assessment of gene regulation and proposes the PB-ERA system as a powerful tool to understand the genetic contribution to common diseases.

## Results

### Fine-tuning enhancer-reporter assays to scale in vivo enhancer detection

ERAs have been the benchmark for the interrogation of the human genome and the study of the effect of variations in the human DNA sequence (4, 51). Standard assessment of enhancer activity in mice is achieved after pronuclear microinjection of zygotes with linearized constructs. ERAs rely on the random integration of the constructs into the genome, which is very inefficient, thus being the main bottleneck in these studies. In addition, the increasing number of predicted enhancers and polymorphisms associated with disease phenotypes urges for the improvement of ERA performance to reduce time, costs and number of animals (28). In order to overcome this limitation, we developed the PB-ERA system, an enhancer assay assisted by transposition. The PB-ERA system relies on the ability of the PB transposase to recognize PB-specific inverted terminal repeats (PBRs) and integrate the inner DNA content into the genome. We microinjected the pPB-βlacZ vector, consisting of the transgene (β-globin minimal core promoter, lacZ reporter gene and candidate genomic region) surrounded by PBRs, in an episomal way together with the mRNA of a hyperactive version of PB (Figure 1A).

**Figure 1.**
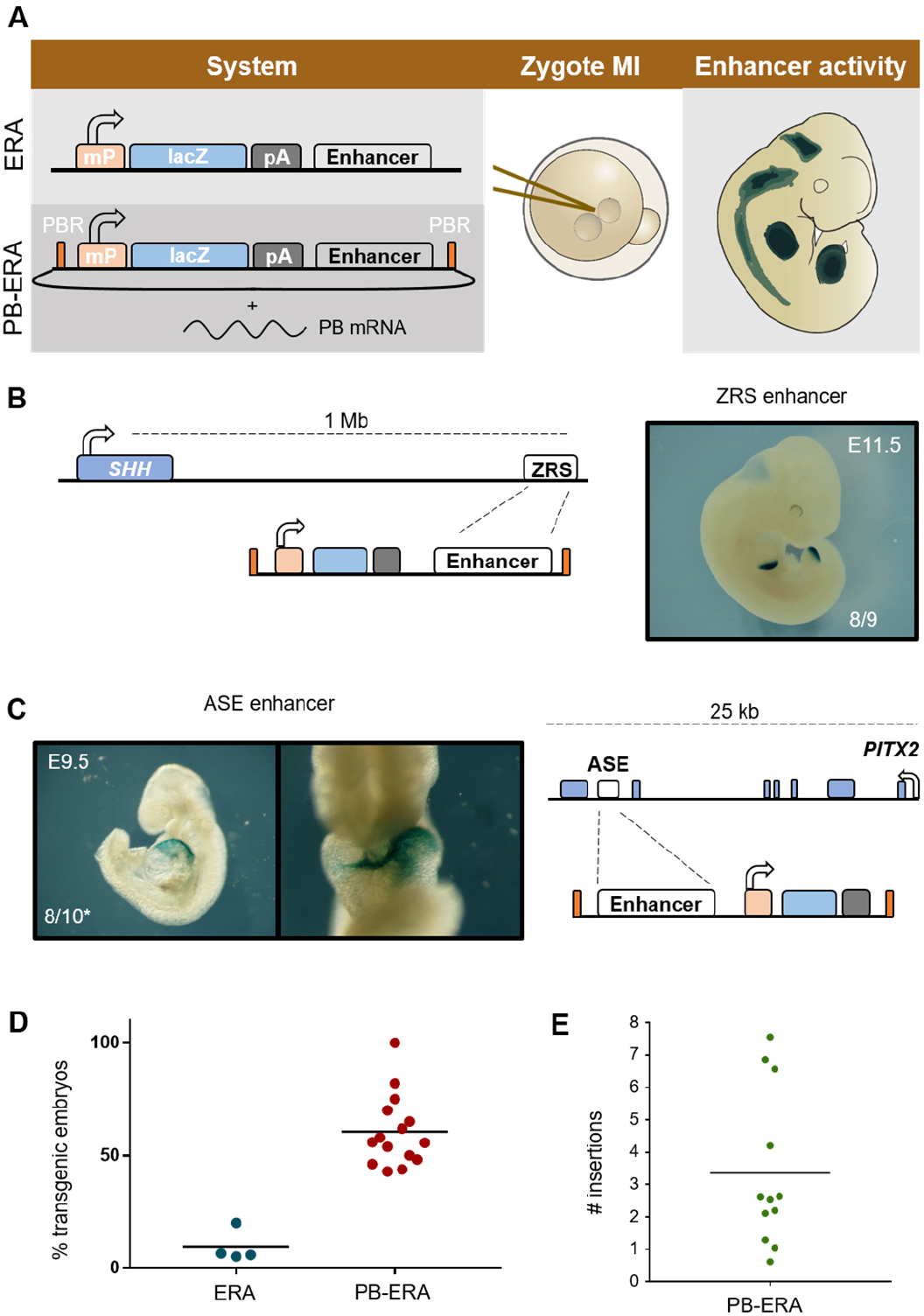
The PB-ERA system is an efficient tool to identify enhancers. **A)** Schematics showing the ERA (linear) and PB-ERA (episomal) vectors that are microinjected into mouse zygotes, to be then transferred to a foster mother until embryos reach the developmental stage of interest (E9.5 E14.5), when they are stained for β-galactosidase activity as a readout of enhancer activity. **B-C)** The PB-ERA system recapitulates the tissue-specific patterns driven by previously characterized human enhancers, such as the ZRS limb enhancer from the *SHH* gene (**B**) or the ASE cardiac enhancer from *PITX2* (**C**). Indicated are the number of lacZ positive embryos versus total transgenic embryos; asterisk in C indicates an estimation of the number of transgenic embryos based on the overall efficiency of the PB-ERA system. **D)** Comparison of the efficiency of transgenesis using classical linearized injection of reporter constructs (ERA) versus the PB-ERA system, as the percentage of transgenic embryos as identified by genotyping versus the total number of embryos obtained. Each data point represents a candidate fragment tested for which several sessions of microinjection where carried out. **E)** Number of integrations in single embryos calculated by qPCR of genomic DNA and normalized to a mouse line with a single copy of the transgene. MI, microinjection; mP, minimal promoter; pA, poly-adenylation signal; PBR, piggyBac repeats.

We first checked the system’s ability to translate the PB mRNA and integrate the transgene early after zygote microinjection (52). To do so, we used GFP driven by the constitutive CAG promoter surrounded by PBRs (pPB-CAG-GFP) and confirmed that at embryonic day (E) 10 embryos were completely fluorescent (Supplementary Figure S1A). This data sustained that transposase-mediated transgenesis could be used to evaluate reporter expression in transient.

Next, we assessed the suitability of the PB-ERA system to capture enhancer activity. To do so, we cloned known enhancers in the pPB-βlacZ vector and tested them by transgenesis. We tested a 2.2kb genomic region containing the human ZRS enhancer, a strong enhancer of *SHH* that is active in the posterior limb bud and has been associated to polydactyly (53, 54). We also tested a 600 bp intronic fragment containing the previously described human left asymmetric enhancer (ASE) of the cardiac transcription factor *PITX2* that drives expression in the developing heart (55).

We saw that both the limb-specific ZRS enhancer and the cardiac ASE enhancer recapitulated the described patterns they drive (Figure 1B, C). It was noteworthy that nearly all generated transgenic embryos, as detected by genotyping, showed β-galactosidase activity (8 out of 9 for the ZRS enhancer; Figure 1B, Supplementary Table S5). To compare if PB_ERA system improves the classical transgenesis we compare the percentage of transgenic rate in all regions tested (15 regions of up to 14,4Kb; Supplementary Table S5) achieving a 60.52% of transgenesis in more than 150 embryos (E9.5-E14.5) as compared to a ~9% of efficiency using classical transgenesis (Figure 1D). Therefore, the PB-ERA system improves conventional transgenesis by more than 6-fold and allows the assessment of enhancer activity in mouse embryos.

Nevertheless, we also found that this system led to a significant number of stained embryos showing no specific pattern when using an empty vector (Supplementary Figure S1B). We reasoned that a large number of genomic insertions, what could potentially be boosting reporter signal unspecifically due to multiple copies, might accompany PB-mediated transgenesis. To verify so, we carried out quantitative genotyping to estimate the number of copies of the construct per embryo, and found an average of 3.4 insertions per transgenic embryo (Figure 1D), a rather low number. We can therefore rule out this as a problem of the system.

### Systematic assessment of enhancer activity in tissue culture finds regulatory elements in AF risk loci

The increased output achieved by the PB-ERA system would allow us to undertake a more systematic evaluation of the regulator activity of several genomic regions. We decided to do so by dissecting ten loci associated with AF to functionally characterize the origin of these associations. We selected 5 kb surrounding the AF risk-associated SNPs for the nine strongest AF-loci detected by GWAS and the intronic CNV (4.3 kb) detected at the 5q35 locus containing the KCNIP1 gene (Table 1). For each of the ten loci, we used the following criteria as predictors of regulatory activity: 1) presence of expression quantitative trait loci (eQTL) for a gene localized within the same topologically associated domain (TAD)(56); 2) histone post-translational modifications associated with active enhancers, such as H3K4me1 or H3K27ac (57, 58); and 3) binding by cardiac transcription factors (GATA4, TBX5, NKX2-5) as detected by ChIP-seq (47, 48). This first categorization showed that all of the selected loci were positive for at least one of the three predictors of enhancers, only with the exception of the *KCNIP1* locus, which suggests that many of the candidate regions might potentially be enhancers (Figure 2A). According to these predictors, three loci stood out from the rest as likely to contain disease-related enhancers: *CAV1*-AF, *C9orf3*-AF and *SYNE2*-AF (Figure 2A) since they fulfill more than two criteria.

**Figure 2.**
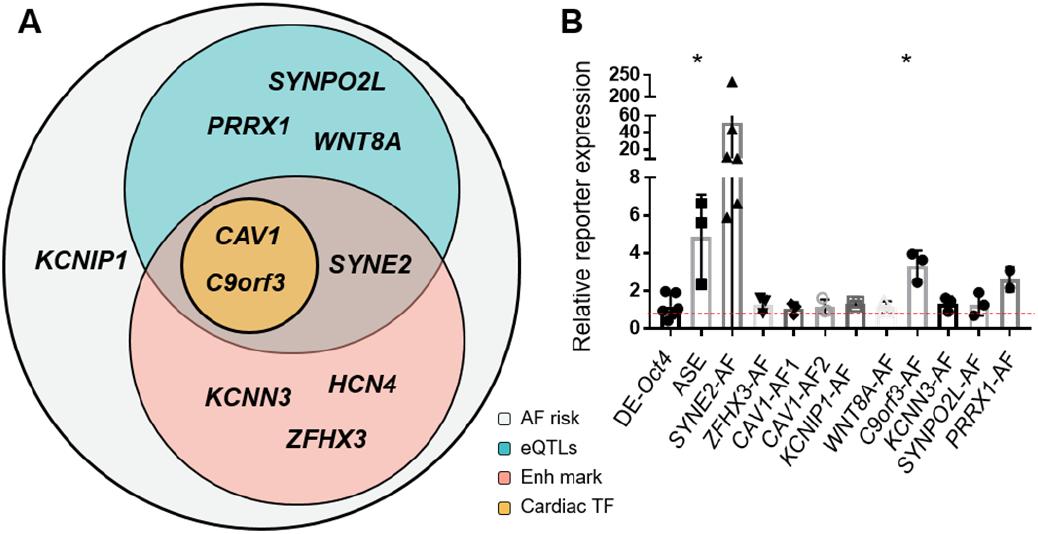
The *SYNE2* and *C9orf3* loci contain potential heart enhancers. **A)** Schematic representation of the regulatory features of the ten AF-associated loci included in this study (grey circle). The three categories included in the Venn diagram are the presence of eQTLs for heart expression of any of the genes in the locus (blue circle), histone marks for active enhancers (H3K4me1 and/or H3K27Ac; red circle), or binding as detected by ChIP-seq peak of cardiac transcription factors (TBX5, GATA4 or NKX2-5) in human or mouse differentiated cardiomyocytes (oranges circle). **B)** Enhancer activity of the candidate loci represented as RNA/DNA ratio after transfection of PB-ERA constructs in the mouse HL-1 cardiac cell line. The distal enhancer (DE) of *Oct4* is used as a negative control and the *PITX2* ASE enhancer as a positive control. Red dashed line represents the mean value of negative control. * p-value < 0.05 compared to the negative control by Student’s t-test.

**Table 1.** Genomic loci analyzed in this study.

| locus | gene/s | tested region (hg19) | variant type | SNPs within region | reference |
| --- | --- | --- | --- | --- | --- |
| 16q22.3 | ZFHX3 | chr16:73049120-73054120 | SNV | rs2106261; rs2359171 | Benjamin et al. 2009 |
| 1q21.3 | KCNN3-PMVK | chr1:154811768-154816768 | SNV | rs6666258; rs13376333 | Ellinor et al. 2010 |
| 1q24.2 | PRRX1-METTL11B | chr1:170566817-170571817 | SNV | rs3903239 | Ellinor et al. 2012 |
| 5q31.2 | WNT8A-FAM13B | chr5:137417489-137422489 | SNV | rs2040862 | Ellinor et al. 2012 |
| 7q31.2 | CAV1-CAV2 | chr7:116183741-116188741 | SNV | rs3807989 | Ellinor et al. 2012 |
| 7q31.2 | CAV1-CAV2 | chr7:116188801-116193801 | SNV | rs11773845 | Ellinor et al. 2012 |
| 9q22 | C9orf3 | chr9:97710959-97715959 | SNV | rs10821415 | Ellinor et al. 2012 |
| 10q22.2 | SYNPO2L-MYOZ1 | chr10:75419016-75423327 | SNV | rs10824026; rs6480708 | Ellinor et al. 2012 |
| 14q23.2 | SYNE2 | chr14:64678348-64683348 | SNV | rs1152591; rs2738413 | Ellinor et al. 2012 |
| 15q24.1 | HCN4 | chr15:73649674-73654674 | SNV | rs7164883 | Ellinor et al. 2012 |
| 5q35.1 | KCNIP1 | chr5:170128728-170132986 | CNV | - | Tsai et al. 2016 |

We cloned the human sequence of all ten selected candidates into the pPB-βlacZ vector in order to functionally assess their enhancer activity. In spite of the limitation of testing in-vitro the activity of putative enhancer that can be active in a very cell-specific context, as a first approach, we transfected the constructs into HL-1 mouse atrial cardiomyocytes (59). This tissue culture model recapitulates general electrophysiological features of cardiomyocytes and the expression of key genes coding for ion channels, gap junction components and cardiac transcription factors, including *Tbx5*, *Gata4*, *NKX2-5* and the atrial-specific *Pitx2* gene (60–62). HL-1 cells were co-transfected with the pPBase vector that expresses PB to enable transposition. As a positive control, we used the same ASE enhancer of *PITX2* that we previously tested in embryos (Figure 1C); for negative control, we used the pPB-βlacZ reporter construct containing the pluripotent-specific *Oct4* distal enhancer (DE) specifically active in mouse pluripotent stem cells (63). We found that two of the tested constructs activated expression in HL-1 cardiomyocytes: *SYNE2*-AF that contains the AF risk-associated SNPs rs2738413 and rs1152591, and *C9orf3*-AF that contains rs10821415. The positive control (ASE) also increased significantly expression (Figure 2B, Table 1). Interestingly, these two genomic regions qualified as very likely to contain enhancers in our analysis above (Figure 2A). This data shows that at least two genomic regions associated with AF risk contain regulatory activity in cardiomyocytes in-vitro.

### The PB-ERA system identifies in vivo enhancers that are not detected in tissue culture assays

We next used the PB-ERA system to test for enhancer activity of the ten selected AF-associated by means of transient mouse embryo transgenesis. To do so, we dissected and examined transgenic embryos at stage E11.5, when the four-chamber heart is already formed and functional, therefore having a higher probability to capture regulatory activity related to AF. Of these ten fragments (Table 1), five did not drive reporter expression in a consistent pattern different to that of the empty vector.

We tested a 4.3 kb fragment located in the first intron of the *KCNIP1* gene (Figure 3A), which has been described as a copy number variation (CNV) associated to AF (64). This fragment acted as a cardiac enhancer, driving reporter expression in scattered cells throughout the heart (Figure 3B, B’). This was rather unexpected, as the *KCNIP1*-AF region did not show any of the predictors for enhancer activity we previously used (Figure 2A). The intron where this enhancer is located interacts with the promoter of the *KCNIP1* gene in human induced pluripotent stem cell-derived cardiomyocytes (hiPSC-CMs), as detected by promoter-capture Hi-C (Supplementary Figure S2A)(50). Indeed, *KCNIP1* is expressed in all four chambers of the heart (64), similar to the enhancer activity that we detected in our transgenic embryos (Figure 3B, B’). Additionally, the intronic region within which this enhancer is located also interacts with the promoter of the *TLX3* gene that is located ~600 kb downstream of the enhancer (Supplementary Figure S2A). Interestingly, TLX3 has been shown to regulate the expression of miR-125b, a microRNA which is critical for fibroblast-to-myofibroblast transition and cardiac fibrosis (65, 66). Further studies will be needed to address a possible role of *TLX3* in cardiac development and homeostasis and in the pathophysiology of AF. Both *C9orf3*-AF and *SYNE2*-AF candidate genomic regions (Figure 3C, E) showed cardiac-enhancer activity in mouse embryos (Figure 3D, F) as well as in cell culture (Figure 2B). The intronic *C9orf3*-AF fragment (Figure 3C) showed only mild expression of the reporter in the heart (Figure 3D), but the risk associated allele at rs10821415 correlated with lower expression of *C9orf3* in atrial tissue, indicating that it might be the target gene of this cardiac enhancer (Supplementary Figure S2B; GTEx data from (44)). *SYNE2*-AF was active in the outflow tract (OFT) and in the lungs of both E11.5 and E14.5 embryos (Figure 3F, Supplementary Figure S2C, D). In the human atria, the expression of *SYNE2* correlated with the genotype of both rs2738413 (Figure 3H) and rs1152591 (Supplementary Figure S2E) variants and it is precisely in the atria where the cardiac expression of the mouse *Syne2* gene is higher (Supplementary Figure S2F; RNA-seq data from 3D-Cardiomics (67)). However, these AF-associated SNPs also correlated with the expression of the neighboring genes (*ESR2* and *MTHFD1*) in the lungs (Supplementary Figure S3B), where this region also showed enhancer activity, suggesting a more complex regulatory activity of this region. *SYNE2* is a large gene that produces two different transcripts (Figure 3E). Interestingly, the *SYNE2*-AF region overlaps the start of the shorter *SYNE2* isoform and the two SNPs associated to AF in this region lie upstream in close proximity to its transcription start site (Figure 3E). Exploring the expression data of the two *SYNE2* transcripts in GTEx (44), we observed that the short isoform is predominantly expressed in skeletal muscle and in the heart, and the long isoform has a lower expression in these tissues (Figure 3G)Although *SYNE2*-AF might contain an alternative promoter for the small muscle isoform of *SYNE2*, that putative promoter activity would be independent from the enhancer activity described here, due to the cloning strategy that we followed in which the insertion site in the pPB-βlacZ vector is located downstream of lacZ. Therefore, the *SYNE2*-AF enhancer harbors cis-regulatory potential capable of triggering reporter expression from downstream. Indeed, if we check promoter-capture Hi-C data in hiPSC-CMs (50) we can see long-range interactions between the *SYNE2*-AF region and the promoter of the large isoform of *SYNE2*, as well as with the promoter of *ESR2* gene (Supplementary Figure S3A).

**Figure 3.**
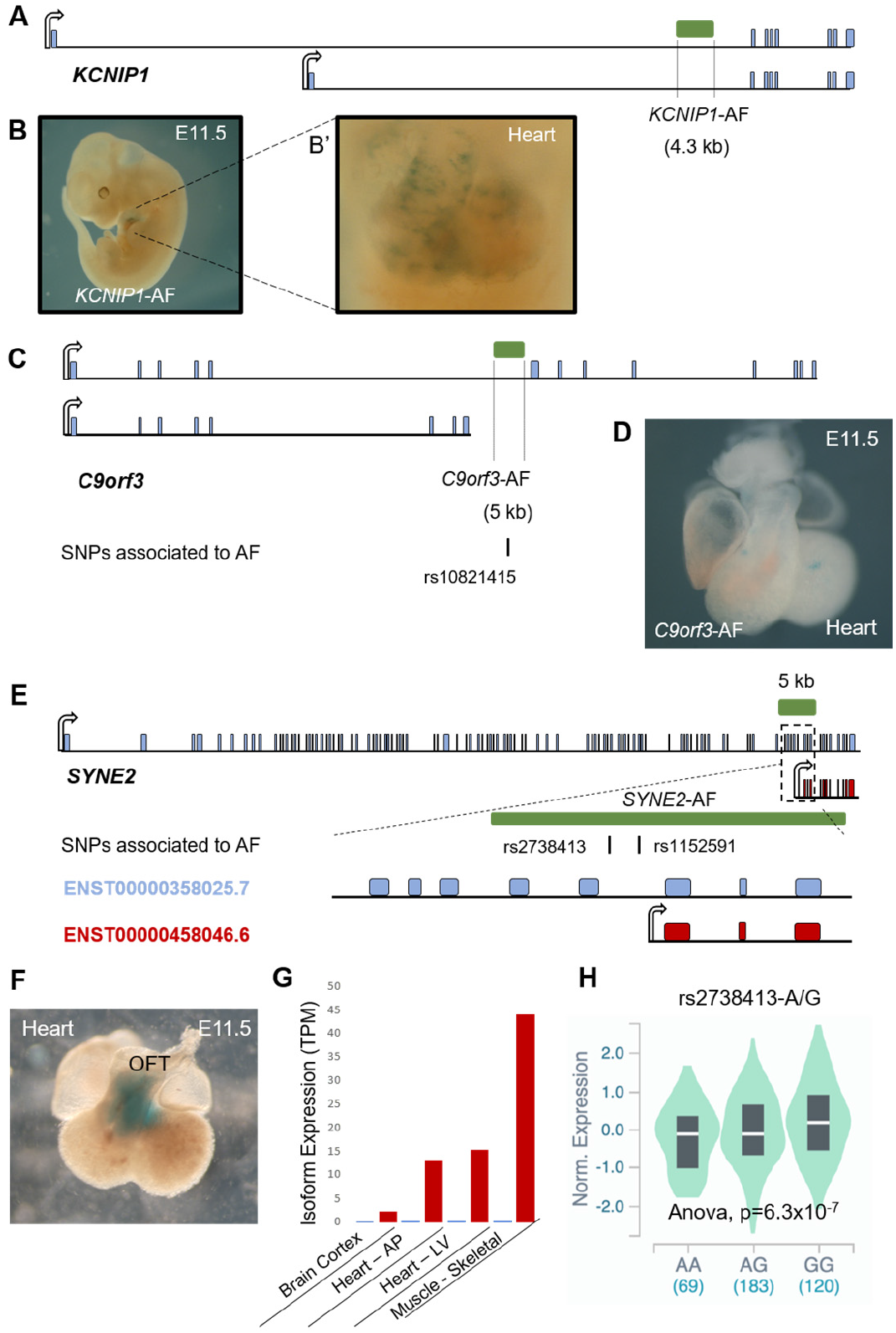
Identification of AF enhancers in vivo. **A)** Schematic representation of the *KCNIP1* locus, showing the location of the *KCNIP1*-AF region in the first intron and 350 kb downstream the promoter. **B)** Cardiac-specific activity of the *KCNIP1*-AF enhancer in E11.5 transgenic embryos with expression in the four chambers of the heart (**B’**). **C)** Schematic representation showing the *C9orf3*-AF region, located in the large intron of the *C9orf3* gene, that contains the AF-associated variant rs10821415. **D)** Dissected heart of an E11.5 transgenic embryo for *C9orf3*-AF. **E)** The *SYNE2* gene encodes a larger (blue) and a shorter (red) isoforms. AF variants rs2738413 and rs1152591 overlap the promoter region of the small isoform and are located in the *SYNE2*-AF fragment. **E)** Enhancer activity for the *SYNE2*-AF fragment in the outflow tract (OFT) of the heart of an E11.5 transgenic embryo. **G)** Expression data from GTEx of the small *SYNE2* isoform in heart and muscle. **H)** Risk A allele of variant rs2738413 correlates with lower *SYNE2* gene expression in atrial tissue (44).

We confirmed the promoter-capture HiC data by circularized chromosome conformation capture (4C-seq) to explore the three-dimensional interactome of this enhancer at an increased resolution. Because we aimed to perform the assays in the physiological context of the heart, we selected the mouse genome region syntenic to human *SYNE2*-AF and analyzed chromatin from the atria of adult mouse hearts. This assay confirmed that the enhancer identified at this locus is interacting, upstream, with *SYNE2* and, downstream, with the promoter of *ESR2* (Supplementary Figure S3C). However, we could not capture interactions between the *SYNE2*-AF enhancer and the promoter of MTHFD1 in adult hearts. Altogether, this data showed that *SYNE2*-AF is a human enhancer regulating the *ESR2* and *MTHFD1* genes in the lung and the *SYNE2* gene in the atria, from which the link with AF might potentially come.

### A complex regulatory landscape of the CAV1 locus suggests novel genes involved in AF

We next selected two genomic regions of 5 kb from the large second intron of *CAV1* for the assessment of enhancer activity in embryos (Figure 4A). These regions contain variants rs3807989 and rs1173845 that are among the strongest GWAS-SNPs associated to AF (23) as well as other variants associated to electrophysiological traits (68). Analysis of available epigenomic data shows the presence of enhancer marks (H3K4me1 and H3K27ac) overlapping these regions in human samples from adult left ventricle (Supplementary Figure S4A). When tested using the PB-ERA system, *CAV1*-AF1 and *CAV1*-AF2 drive reporter expression in the heart of transgenic embryos (Figure 4B, B’, C and C’). Indeed, *CAV1*-AF1 is bound by cardiac transcription factors (GATA4 and TBX5) in human differentiated cardiomyocytes (Supplementary Figure S4B; data from (47)) as well as the orthologous mouse region, which is bound by GATA4, TBX5 and NKX2-5, in mouse differentiated cardiomyocytes (Supplementary Figure S4C; data from (48)). Together, this data shows that *CAV1*-AF1 and *CAV1*-AF2 are regulatory elements active in the heart, whose function is very likely conserved in mammals. *CAV1* encodes Caveolin-1, the main component of caveolae which are small membrane invaginations present in many cells, including cardiomyocytes (69). Caveolin-1 might play a relevant role in the electrophysiological and mechanical properties of cardiomyocytes interacting with ion channels (70) and gap junction proteins (71) and since caveolae regulate plasma membrane curvature which prevents membrane rupture (72). Accordingly, knockout mice for *Cav1* have cardiac conduction affected and develop ventricular arrhythmias (73). The identification of cardiac enhancers at the AF-risk locus might help elucidate the mechanism behind the association. However, there is no direct evidence that these enhancers control the expression of *CAV1* or any other gene within the TAD.

**Figure 4.**
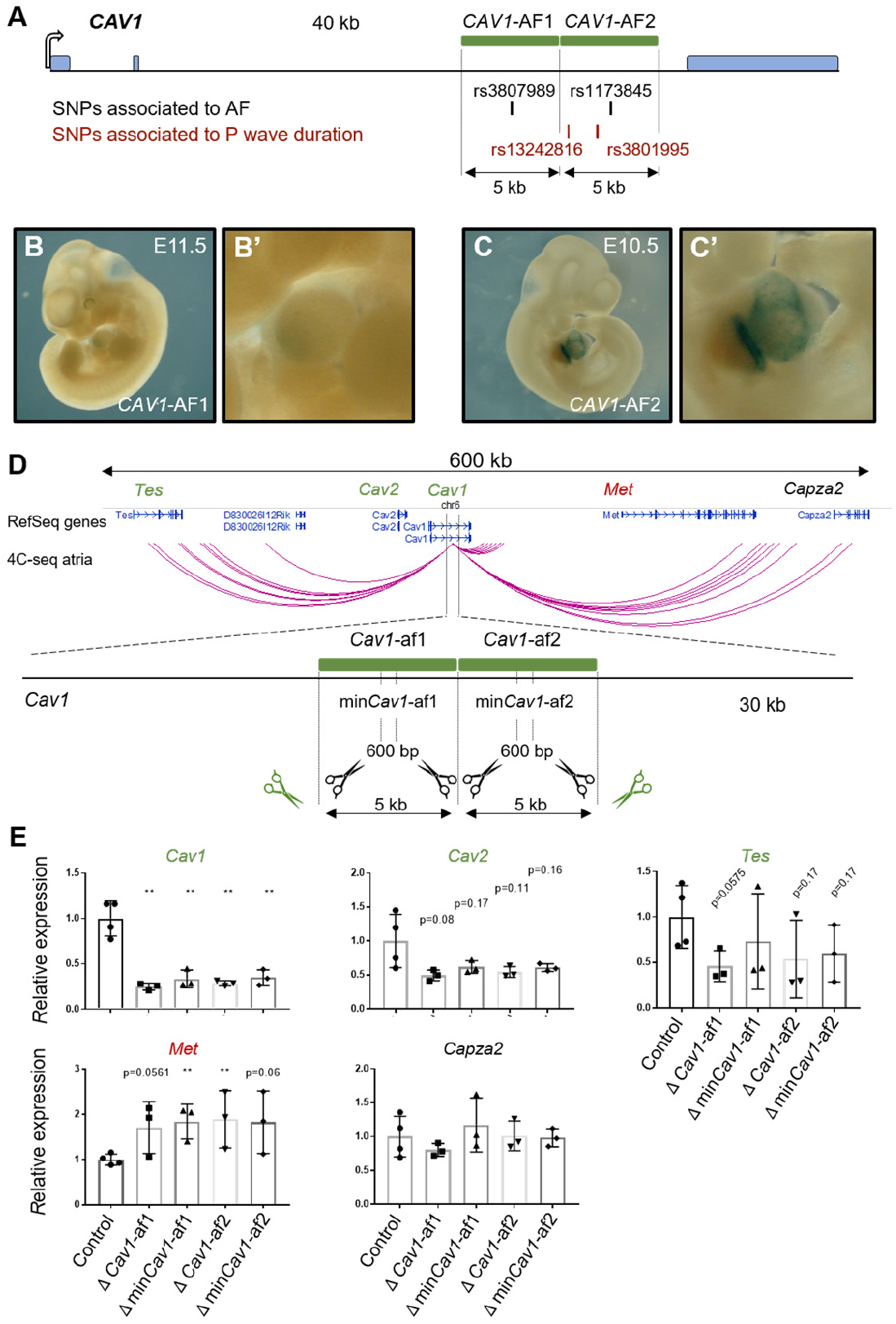
The 7q31 AF risk locus contains cardiac regulatory elements differentially regulating upstream and downstream genes. **A)** Schematic representation of the *CAV1* gene, indicating the candidate enhancer regions (*CAV1*-AF1 and *CAV1*-AF2; green boxes) and SNPs associated to AF (black) or electrophysiological traits (P wave duration; red). **B-C)** Enhancer activity in the heart of transgenic embryos for the *CAV1*-AF1 (**B** and **B’**) and the *CAV1*-AF2 (**C** and **C’**) regions. **D)** 4C-seq in the atria of mouse hearts showing significant interactions, where the orthologous region of *CAV1*-AF1 and *CAV1*-AF1 interact with multiple promoters and gene bodies within the locus. A schematic representation of the large intron of *Cav1* in the mouse is depicted below, indicating the targeted deletions of the orthologous candidate enhancer regions. **E)** Deletions of *Cav1*-af1 and *Cav1*-af2 in HL-1 cardiac cells led to downregulation of upstream genes (*Cav1*, *Cav2* and *Tes*), and to upregulation of the downstream *Met* gene, while not affecting *Capza2*. ** p-value < 0.01 compared to the control by Student’s t-test. Other p-values > 0,05 are indicated above each graph.

The 900 kb of the 7q31 locus are highly interconnected as shown by promoter-capture Hi-C data in differentiated cardiomyocytes from hiPSC (50). Within this locus, the large intron of *CAV1* that contains the regulatory elements that we identified is involved in long-range chromatin interactions (Supplementary Figure S4A). As both the sequence and epigenetic features of the enhancers located in the second intron of the human *CAV1* were conserved in mammals, we performed 4C-seq of the orthologous enhancer region in the atria of adult mouse hearts. This analysis showed that the enhancers interacted long range with other genes within the TAD containing *Cav1*, such as *Tes* (~300 kb upstream), *Met* (~100 kb downstream) and *Capza2* (~300 kb downstream)(Figure 4D). Due to proximity, we could not use the 4C data to capture interactions with the promoters of *Cav1* and *Cav2*. However, the general three-dimensional landscape of the human region between *CAV1* and *CAV2* showed that this region widely interacts with each other (Supplementary Figure S4A). Altogether, this data indicates that the cardiac regulatory elements identified in the second intron of *CAV1* might regulate multiple genes within a region of 700 kb.

### AF-associated regulatory elements at the CAV1 locus differentially regulate upstream and downstream genes

Enhancer-gene interaction is not sufficient to determine direct gene regulation (74). In order to identify the target genes whose expression is regulated by *CAV1*-AF1 and *CAV1*-AF2, we used the CRISPR/Cas9 system to disrupt the orthologous regions to these CREs in mouse HL-1 atrial-like cardiomyocytes and then examined gene expression by RT-qPCR. We designed pairs of sgRNAs to delete each 5 kb enhancer separately (*Cav1*-af1, *Cav1*-af2; Figure 4D). The deletion of either regulatory element led to significant downregulation of *Cav1*, and a clear trend to decrease *Cav2* and *Tes* expression (Figure 4E), confirming that *Cav1*-af1 and *Cav1*-af2 are acting as enhancers of these three genes located upstream. Surprisingly, *Cav1*-af1 and *Cav1*-af2 seemed to be negatively regulating the downstream gene *Met* (Figure 4E). The interaction between the heart enhancers with the *Capza2* gene did not affect gene expression in this cell type (Figure 4E). Next, we wanted to investigate whether the variants rs3807989 and rs1173845 located within *CAV1*-AF1 and *CAV1*-AF2, respectively, were at an essential position in the CREs. To do so, we designed pairs of sgRNAs targeting minimal regions of ~600 bp (min*Cav1*-af1 and min*Cav1*-af2) surrounding the orthologous region in mouse to that in human containing the AF-risk associated SNPs. Deletion of any of the core regions led to comparable levels of downregulation of *Cav1*, *Cav2* and *Tes*, and upregulation of *Met*, in HL-1 cells to that of the 5 kb genomic fragments (Figure 4E). Overall, our data suggest that *CAV1*-AF1 and *CAV1*-AF2 are highly conserved heart regulatory elements that contain AF variants at core domains essential for their function and act differentially on genes located either upstream or downstream.

### The PB-ERA system identifies a negative regulator associated to AF in the first intron of ZFHX3

Variants within the first intron of *ZFHX3* at the 16q22 locus have been associated to AF since the very first genetic reports more than a decade ago (20, 21), but we still lack functional evidence that *ZFHX3* or other genes are regulated by the 16q22 AF-risk locus (75). To fill this gap, we investigated the regulatory role of a 5 kb region, namely *ZFHX3*-AF, in the first intron of *ZFHX3* that contains two variants associated with AF, rs2106261 and rs2359171 (Figure 5A), by testing its regulatory activity using the PB-ERA system in E11.5.transgenic embryos.

**Figure 5.**
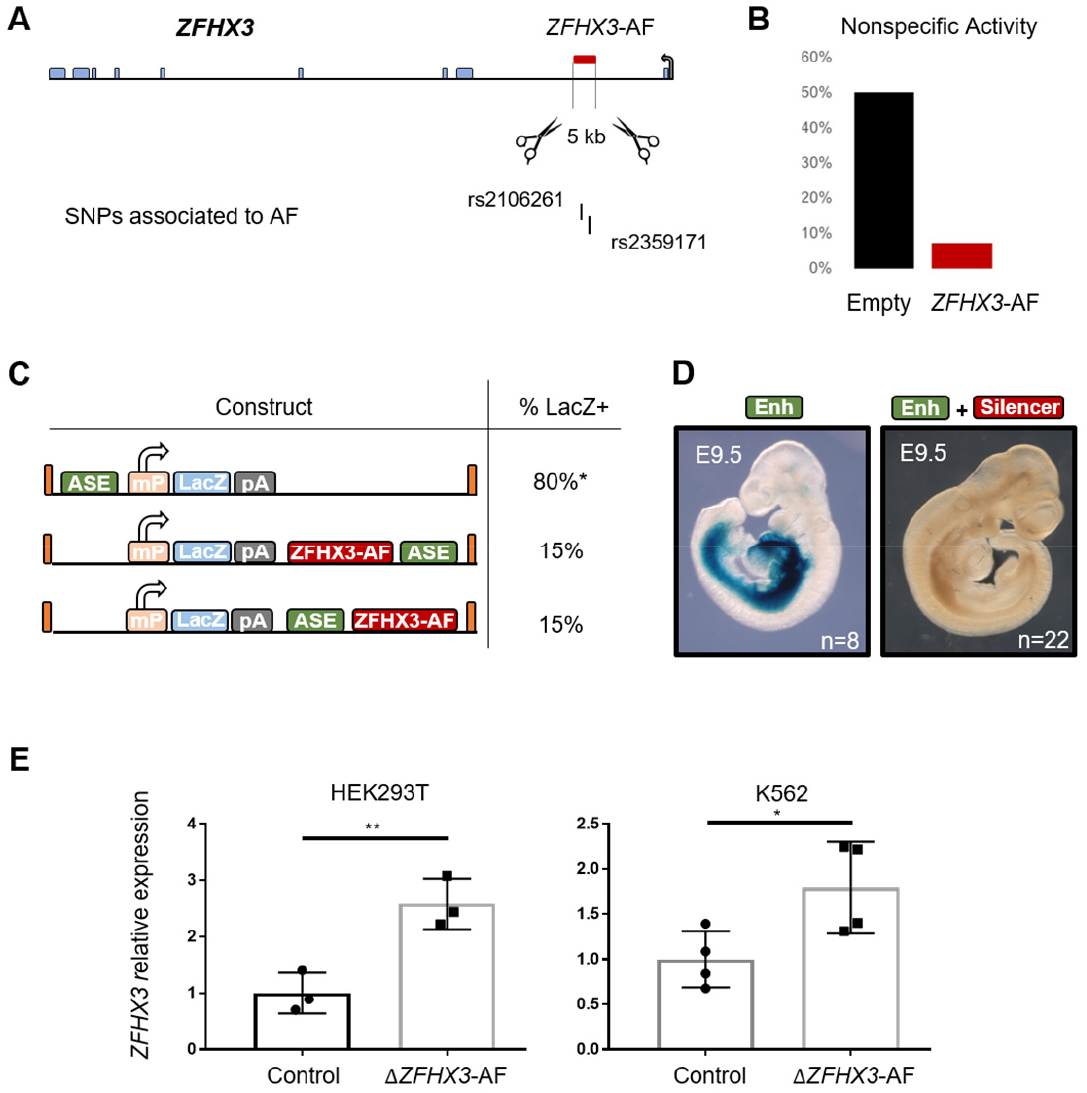
The candidate *ZFHX3*-AF region is a ubiquitous negative regulator. **A)** Schematic representation of the *ZFHX3* gene indicating the region containing the risk variants rs2106261 and rs2359171 (*ZFHX3*-AF, red box). **B)** Percentage of transgenic embryos showing unspecific reporter activity with the empty PB-ERA vector or with the same construct where the *ZFHX3*-AF region is cloned. **C)** Schematic representation of the chimeric and control constructs tested. Asterisk indicates the estimated proportion of *lacZ* positive embryos (80%) for the ASE element, as embryos could not be genotyped. **D)** Representative embryos showing reporter expression driven by the ASE enhancer (left) or complete silencing when the *ZFHX3*-AF element is cloned together with the ASE (right). **E)** Deletion of the *ZFHX3*-AF element resulted in overexpression of *ZFHX3* in the human cell lines HEK293T and K562. * p-value < 0.05, ** p-value < 0.01, compared to the control by Student’s t-test.

Unexpectedly, the *ZFHX3*-AF genomic region dramatically reduced nonspecific activity as compared to the empty pPB-βlacZ vector (Figure 5B). It should be noted that the high efficiency of transgenesis using the PB-ERA system led to unspecific *lacZ* expression and β-galactosidase activity in up to 50% of embryos, as we observed when no genomic fragment was placed in the construct and we generated transgenics with the empty vector. Noise was easily discriminated from signal due to absence of reproducibility or lower intensity than bona-fide enhancer-driven β-galactosidase activity (Figure S1B). A possible interpretation of this observation would be that this region was acting as a repressive element.

To test this hypothesis further, we used the PB-ERA system to perform *in vivo* assays of the putative enhancer blocking activity of *ZFHX3*-AF on the *PITX2* cardiac ASE (see Figure 1E). We examined E9.5 transgenic embryos carrying one of the following three constructs: the ASE alone (pPB-ASE-βlacZ), the *ZFHX3*-AF located between the βglobin promoter and the ASE (pPB-βlacZ-ZFHX3AF-ASE) or the *ZFHX3*-AF located after the βglobin promoter and the ASE (pPB-βlacZ-ASE-ZFHX3AF) (Figure 5C). We found that both of the two configurations tested, the *ZFHX3*-AF fragment reduced ASE activity independently from its relative position to the promoter and enhancer (Figure 5C, D; Supplementary Table S5). This result, together with the overall reduction of non-specific reporter activity described above, indicates that this region is not acting as an insulator (that would have been positon-dependent), but rather as a silencer in the developing heart.

Next, we analyzed the effect of *ZFHX3*-AF deletion in human cells. We carried out CRISPR/Cas9 genome editing of the region in two human cell lines of different origin, the adherent embryonic kidney HEK293T cell line and the lymphoblastic myeloid K562 cell line. Consistently, the deletion of the AF-associated negative regulator led to upregulation of *ZFHX3* in both cell lines (Figure 5E). This result confirms that the *ZFHX3*-AF fragment is a silencer, which acts in different cell types.

## Discussion

Precise control of gene expression is essential for the correct development and functioning of tissues and organs (12, 76, 77). Therefore, identifying and characterizing regulatory elements and how genetic variation affects their activity is crucial towards achieving precision medicine. In this work, we aimed to improve the available tools for the functional characterization of regulatory elements, and use them to elucidate the genetic mechanism behind the most significant GWAS associations for AF, a major cardiovascular arrhythmia that affects over 30 million people worldwide (17).

After the discovery of the first viral enhancer (78, 79), classical transgenesis allowed for the *in vivo* characterization of mammalian enhancers (6–8). While mouse transgenesis has effectively identified hundreds of regulatory elements involved in mammalian development and disease (3, 5, 51, 53, 80), methods of *in vivo* ERAs are rather inefficient (81, 82) and have very limitedly evolved in forty years (83, 84). Whereas transposases, such as the Tol2 system (85), have increased ERA performance in zebrafish (85–87), mice were not suitable for rapid and higher scale experiments (87). The Sleeping Beauty (88) or *piggyBac* (33) systems have been used to generate mouse lines with integrated sensors of enhancer activity (89–93). However, segregation of the multiple insertions and enhancer analysis ultimately requires high periods and costs of animal breeding and maintenance.

Our findings that the PB-ERA system is a convenient method to systematically assess enhancer activity via transient transgenesis in disease-associated genomic regions, opens the door to in-depth interrogation of the ever-increasing number of risk loci. Recently, another study aiming to scale up mouse ERAs developed 3-component method assisted by CRISPR in order to test the impact of human variants in the ZRS enhancer (94). This, once again, stresses the need for mouse *in vivo* ERAs of higher throughput. Here, we showed that the 2-component PB-ERA system yields an average 60% of transgenic embryos, being the most efficient system reported to the best of our knowledge. The number of integrations per transgenic embryo were not high, thus enabling the capture of enhancer patterns with a minimized position effect. Furthermore, and due to an intrinsically high non-specific background, the PB-ERA system that we implemented was able to detect genomic regions with both enhancer and repressing activity, being able to discriminate between silencers and insulators, and overcoming an existing limitation in the field that mostly focused on positive regulators of gene expression.

In the context of AF, insights into how GWAS associations contribute to the disease remains a challenge even for the most significant SNPs at the 4q25 locus. AF-SNPs in this locus lie within a gene desert where we and others found distal enhancer elements that interact with the promoters of *PITX2* and *ENPEP* (34, 95, 96). On the one hand, *PITX2* encodes a developmental TF expressed in the atria (97) found to be decreased in AF patients (98, 99) and in a sheep model of induced AF (100). On the other hand, *ENPEP* encodes aminopeptidase A, a member of the reninangiotensin system that cleaves angiotensin II to produce angiotensin III and is involved in hypertension (101). Nevertheless, the variants in the locus do not correlate with the atrial expression of *PITX2* in patients (102), illustrating how complex it is to ascertain the contribution of human variation to complex diseases. Interestingly, some of the regulatory elements in this locus showed non-tissue-specific potentiator activity (34). This further supports the idea that some GWAS variants do not necessarily disrupt the activity of canonical tissue-specific enhancers, or that there is redundancy of regulatory elements. Any of these options would explain why regulatory GWAS variants suggests might have a mild effect on gene expression.

In the last decade, GWAS performed in over two million people, including 200,000 AF cases (20, 23–27, 103–111), have identified ~130 risk loci, identifying additional genomic regions associated through less characterized forms of genetic variation such as indels and CNVs (13, 112). Of these, our study addresses ten of the most significant associations and assesses their regulatory potential and target genes. While we observe an overlap between enhancers identified in cultured atrial myocytes and mouse transgenic embryos, it is however not surprising that our in vivo approach extends cell culture experiments. Our study shows that prioritization of candidate loci increases the success rate of enhancer identification as we found enhancers in all three human loci presenting three layers of enhancer marks, i.e. 7q31, 9q22 and 14q23. It is important to highlight, though, that we are still far from completely understanding the regulatory genome and cataloguing all CREs. The cardiac enhancer identified at the *KCNIP1* locus is an example of the former, since no other predictor supported this candidate beyond the previous association to AF of a 4.4 kb CNV. In this case, chromatin analysis of the region involved not only *KCNIP1* but also the transcription factor-encoding *TLX3* gene, and it will be very interesting to explore *TLX3* expression in patients and its putative role in AF. On the other hand, half of the tested regions did not show activity, what suggests that these AF-associated genomic regions have other functions not observable by our regulatory assays.

The functional analysis performed at the 7q31 locus indicate that the large second intron of the *CAV1* gene might be a hub of enhancers or a cardiac ‘super-enhancer’ (113). The 10 kb spanning the two regulatory elements that we described (*CAV1*-AF1 and *CAV1*-AF2) might harbor modules that cooperatively regulate gene expression. Furthermore, the 7q31 locus is an example of how important it is to assess enhancer activity in its native chromatin region. Enhancer deletion using CRISPR genome editing not only confirmed that smaller regions containing variants rs3807989 and rs1173845 are critical for gene regulation, but also identified their target genes. These enhancers regulate *CAV1* and *CAV2*, that code for proteins that are members of caveolae and are involved in mechanosensing. More surprising was that they also regulated *MET* and *TES*. *MET* encodes hepatocyte growth factor (HGF) receptor that plays a physiological cardio-protective role in adult cardiomyocytes preventing cardiomyocyte hypertrophy, heart fibrosis, and heart dysfunction (114). On the other hand, *TES* encodes Testin, a member of the focal adhesions that connects the cell to the extracellular matrix and is involved in mechanical and regulatory signal transduction (115, 116). *TES* have been associated to other cardiovascular diseases such as atherosclerosis and aneurism, playing important roles in endothelial and vascular smooth muscle cells where Testin can be found in the nucleus putatively coregulating gene expression (117, 118). Therefore, our analysis has uncovered a putative role of these genes in AF that might be of clinical relevance.

Silencers are also essential in the coordinated regulation of gene expression and while recent reports have developed methods for their scalable identification in cell culture (119, 120), currently there were no mouse in vivo tools for their efficient characterization. The genetic mechanism behind the AF associations at the 16q22 has remained elusive after a decade of research, but here we identify a *ZFHX3*-AF silencer at the 16q22 locus that directly regulates *ZFHX3* gene expression and is able to outcompete heart enhancers *in vivo*. *ZFHX3*, also known as *ATBF1*, encodes a developmental transcription factor that has been involved in myogenic and neuronal differentiation (121, 122). Our findings on the 16q22 would suggest that unregulated expression of *ZFHX3* in cardiac cells could lead to AF. Our work also stresses the importance of other types of cis-regulatory elements beyond enhancers, such as silencers, when understanding the genetic contribution to disease risk, providing a tool for their study.

## Materials and Methods

### Cloning

The pPB-βlacZ vector was obtained by inserting a cassette (3.7 kb) containing a β-globin minimal promoter, a *lacZ* reporter gene and a SV40 polyadenylation signal from the p1230 plasmid (34) into a pPB-CAG-DDdCAs9-VP192-T2A-GFP vector (35), after removal of the CAG-DDdCAs9-VP192-T2A-GFP cassette (digested with SpeI and BamHI). The *lacZ* cassette was amplified using primers ‘PB-Cassette Fw’ and ‘PB-Cassette Rv’ (Supplementary Table S1)

Commercial human DNA (Promega, Cat. No. G1521) was used for PCR amplification of all tested genome fragments from AF associated loci (for primers used see Supplementary Table S1) using Expand High Fidelity PCR system (Roche ref. 11732650001). Primers were design using NEBuilder assembly tool to have 20-basepair (bp) homology arm overhangs for Gibson cloning (36)(NEB, Cat. No. E2611) into the pPB-βlacZ vector digested with SpeI and SacII for 3’ cloning, or HindIII for 5’ cloning. All constructs were verified by Sanger sequencing of plasmids. Large or hard-to-amplify fragments were sub-divided into several PCR fragments with shared homology between them for sequential ligation (colored lower-case nucleotides; Table S1).

Chimeric constructs ASE-ZFHX3 and ZFHX3-ASE were obtained by cloning the ASE fragment upstream or downstream the ZFHX3 fragment, respectively, in the pPB-βlacZ-ZFHX3 vector digested with SpeI (ASE-ZFHX3) or with SacII (ZFHX3-ASE). Specific primers were used in each chimeric design to amplify the ASE fragment with homology to each of the cloning positions.

### Cell culture transfection and CRISPR experiments

Mouse HL-1 atrial cardiomyocytes were cultured in Claycomb medium (Sigma) supplemented with 10% (v/v) inactive (56°C, 30 minutes) fetal bovine serum (FBS) (Sigma), 4 mmol/L L-glutamine (Sigma), 100 μmol/L norepinephrine (Sigma) and 100 U/mL penicillin-streptomycin (Sigma). All seeding supports were previously coated for 24 hours with a solution of gelatin (0.02% w/v, Sigma) and fibronectin (25 μg/mL, Sigma). Human HEK293T embryonic kidney cells were cultured in Dulbecco’s modified Eagle’s medium (DMEM, Sigma) supplemented with 10% FBS, 4 mmol/L L-glutamine and 100 U/mL penicillinstreptomycin. Human K562 erythroleukemia cells were cultured in Roswell Park Memorial Institute (RPMI, Life Technologies) 1640 medium supplemented with 10% FBS, 4 mmol/L L-glutamine and 100 U/mL penicillin-streptomycin.

For PB-ERA experiments, cells were counted one day before transfections and plated at a density of 3 × 10^5^ cells per p12 well (HL-1 cells) or 1.5 × 10^5^ cells per p24 (HEK293T) with complete growth medium. Cells were co-transfected with 1 μg of pPB-βlacZ vector containing the appropriate genomic fragment and 1 μg of the transposase plasmid PBase (a gift from Diego Balboa and Timo Otonkoski). pPB-CAG-GFP was transfected in parallel as an internal control of transfection efficiency. Co-transfections were performed with 6 μl of Lipofectamine 2000 (Invitrogen) diluted in Opti-MEM (Sigma) reduced-serum medium, according to the manufacturer’s protocol. Cells were transferred to complete medium after five hours. The empty vector pPB-βlacZ or the pPB-βlacZ-DEOct4 containing the pluripotent-specific distal enhancer of *Oct4* was used as a negative control. Ninety-six hours after transfection, both DNA and RNA were isolated using AllPrep DNA/RNA kit (Qiagen, Cat. No. 80204) and kept at −80°C for qPCR analysis.

For enhancer deletion in mouse and human cells we transfected the CRISPR/Cas9 gene-editing tool, as described (37). Briefly, 3 x 10^6^ cells were seeded in 10-cm plates the day before transfection. Cells were transfected for five hours with 60 μl of Lipofectamine 2000 and 10 μg of each of the plasmids pSpCas9(BB)-2A-GFP (PX458, Addgene #48138) and pSpCas9(BB)-2A-Puro (PX459, Addgene #48139). Two guides were designed per enhancer (Supplementary Table S2) using CRISPOR (http://crispor.tefor.net/) (38) or Benchling (https://www.benchling.com/) and cloned into either the plasmid containing Cas9-GFP or Cas9-Puro. Forty-eight hours after transfection, GFP+ cells were sorted using Aria Cell Sorter (BD Biosciences) and seeded with puromycin for other four days. Isolated RNA was stored at −80°C for qPCR analysis.

### Quantitative PCR

Isolated RNA was reverse transcribed using the High Capacity cDNA Reverse Transcription Kit (Applied Biosystems). cDNA was used for quantitative PCR (qPCR) with Power SYBR Green (Applied Biosystems) in a 7900HT Fast Real-Time PCR System (Applied Biosystems). Expression of each gene was normalized to the expression of the housekeeping genes *Actin* (HL-1 cells; mouse), *ACTIN* (HEK293T; human) or *GAPDH* (K562; human). Primers used are listed in Supplementary Table S3.

Relative regulatory enhancer activity after PB-ERA assays in cells was calculated as the ratio of reporter *lacZ* expression (RNA) to transfection efficiency (DNA), expressed as mean ± standard deviation, and statistically analyzed by unpaired Student’s t-test (Graphpad Prism5). A minimum of three replicates were used to calculate enhancer activity.

The effect of enhancer deletion was calculated by comparing gene expression of experimental GFP+, Puro resistant cells versus wild type cells transfected with no guide RNAs. A minimum of three replicates were used to assess the effect of enhancer deletion.

### In vitro transcription of the PB transposase

PB transposase was in vitro transcribed from a linear template containing a T7 promoter (T7p) and the cDNA of a hyperactive PB transposase (39). First, linear template was obtained by PCR amplification from the PBase vector, using the primers ‘PB-transcription Fw’ and ‘PB-transcription Rv’ (Supplementary Table S4). Product from PCR amplification (V=50 μl; 1 ul of vector [20 ng/ul]; 1.5 ul each primer [10 uM]; program: 94 °C, 2 min; 10x (94 °C, 15 sec; 65 °C, 30 sec; 72 °C, 2 min); 20x (94 °C, 15 sec; 65 °C, 30 sec; 72 °C, 2 min + 5 sec each cycle); 72 °C, 7 min; 4 °C, ∞) was run in a 1% agarose gel and the desired band (1.8 kb) was purified using QIAquick gel extraction kit (Qiagen, Cat. No. 28704) and used as a template for the transcription reaction. For in vitro transcription, we used ‘mMESSAGE mMACHINE T7 ULTRA Transcription kit’ (Invitrogen, Cat. No. AM1345) at 37 °C for 2 hours, according to manufacturer’s instructions and using 500 ng of template DNA. This kit includes a step to cleave template DNA and polyadenylate the resulting mRNA. The final capped and polyadenylated mRNA was purified using RNA cleanup step from the ‘RNeasy Mini Kit’ (Qiagen Cat. No. 74106) and eluted in 40 μl of nuclease-free water. RNA concentration was measured using Nanodrop (approximate yield: 1.0-1.2 μg/μl) and the product was aliquoted in PCR tubes (2-3 μl each) and stored at −80 °C.

### Transient transgenic assay using the PB-ERA system

For the generation of transient transgenics, F1 (C57Bl/6 x CBA/J) females were superovulated to obtain fertilized oocytes and injected zygotes were transferred to CD1 recipients following standard procedures (40). Episomal non-digested pPB-βlacZ-derived constructs were microinjected into fertilized E0.5 oocytes at a concentration of 2 ng/μl. In vitro transcribed mRNA was microinjected at a concentration of 75 ng/ul together with each pPB-βlacZ construct. To ensure the correct translation of microinjected mRNA, the microinjection needle was aimed at the pronucleus and then removed slowly to ensure that mRNA was partially released in the cytoplasm. A summary of transgenic assays is shown in Supplementary Table S5.

### lacZ staining and genotyping

At the desired stage, pregnant female mice were euthanized and embryos dissected and stained for β-galactosidase activity (40). All embryos were genotyped for *lacZ* by PCR, using primers for *Myogenin* (Supplementary Table S6) as an internal control of a single-copy gene in genomic DNA. Transgenic efficiency was calculated as the percentage of embryos expressing *lacZ* of total transgenics (see Supplementary Table S5).

In order to quantify the number of transgene insertions, we performed qPCR of genomic DNA from transgenic embryos using qPCR primers for lacZ (Supplementary Table S3), and calculated copy number as the relative value to that of heterozygous (one copy of the transgene) and homozygous (2 copies) mouse lines.

### 4C-seq assays and analysis

4C was performed as previously described (41, 42) with minor modifications. Briefly, pools of 20 atria and of 4 ventricles from 8-week C57BL/6 male mice were disaggregated and crosslinked with 2% paraformaldehyde. After fixation, pools were lysed in 50mM Tris-HCl pH7.5, 150 mM NaCl, 5 mM EDTA, 0.5% NP-40, 1% TX-100 and 1x complete protease inhibitor (Roche), digested first with DpnII (New England Biolabs, Cat. No. R0543M) followed by either NlaIII (New England Biolabs, Cat. No. R0125L) or Csp6I (Fermentas, cat.no. ER0211), depending on the initial design of the 4C viewpoint, and re-ligated. For all experiments, 500 ng of the resulting 4C template was used for the subsequent library preparation (primers used are detailed in Supplementary Table S7). Before sequencing, libraries were purified using High Pure PCR Product Purification Kit (Roche), followed by a second purification with Agencourt AMPure Kit to fully remove primers and small fragments. Sequencing was performed by the CNIC Genomics Unit on an Illumina HiSeq 2500 following standard protocols. All sequencing data has been deposited at GEO under accession number XXXXXX. 4C-seq reads were mapped to the MGSCv37 (mm9) mouse reference genome using Bowtie-1.1.2. Reads located to fragments flanked by two restriction sites of the same enzyme, to fragments smaller than 40 bp, or within a window of 10 kilobase (kb) around the viewpoint were filtered out. Mapped reads were then converted to reads per first enzyme fragment ends and smoothened using a 30-fragment mean running window algorithm. Smoothened scores of each experiment were then normalized by the total number or reads.

For contact estimation, the reference genome was digested in silico according to the first and second restriction enzymes. Then, processed reads were assigned to their corresponding first cutter digested genome fragment, filtering out reads on fragments located 5kb around the viewpoint. Quantification was performed considering each fragment end as one captured site if one or more sequences mapped to it. The number of captured sites was summarized per 30 fragments window (max of 60 captured sites per window). The frequency of captured sites per window was used to fit a distance decreasing monotone function and Z-scores were calculated from its residuals using a modified version of FourCSeq (43). Significant contacts were considered in cases where the Z-score was >2 in both replicates and deviated significantly (adjusted p value <0.05) from its normal cumulative distribution in at least one of the replicates. After data analysis, processed reads and interactions were visualized at the WashU EpiGenome Browser Mouse genome (mm9).

### In silico annotation of AF candidate CREs

AF-associated genomic regions were classified according to the presence of regulatory features. For eQTLs, we included AF-SNPs within the candidate fragments and used GTEx publicly available data (44) to annotate them if the expression of any of the genes within the risk locus associated to the genotype of the variants in heart tissue. Histone marks of active enhancers (H3K27ac and H3K4me1) were explored within human candidates and their orthologs in the mouse genome using available data from ENCODE and Roadmap Epigenomics for human left ventricle, right atria and fetal heart, as well as mouse embryonic (E14.5) and adult (8 weeks) heart tissue (45, 46). ChIP-seq data for TBX5, GATA4 and NKX2-5 were used in differentiated cardiomyocytes from both hiPSC (47) and mESC (48) to annotate cardiac TF binding within candidate AF-CREs. Available H3K27me3 ChIP-seq data from human tissues (aorta, left ventricle, fetal heart, fetal kidney, fetal lung, fetal brain, H1 derived neuronal progenitor, H1 ESC, H9 ESC) were used to explore the repressive marks at the *ZFHX3* promoter and AF-associated genomic region. (45, 46). Mouse orthologs of human AF-candidate regions assayed by PB-ERA transfection and transgenesis were obtained using UCSC liftOver tool (49).

Available promoter-capture Hi-C data from hiPSCs and differentiated CM were used to explore putative target promoters interacting with candidate regulatory regions and the specificity of their interaction (50). Tracks were loaded to WashU epigenome browser to represent the data as arcs. For detailed assessment of the overlap between interactions and AF variants, data was represented as the mapping reads of the crosslinked interaction.

### Statistics

Statistical analyses were performed with GraphPad Prism 6 or Microsoft Excel. Data are presented as means ± standard deviation (sd) unless stated otherwise. Asterisks indicate p-values < 0.05. Tests used to calculate p-value are detailed in the figure legends. In general, Student’s t-test was used to compare two groups.

### Animal handling

Mice were housed and maintained in the animal facility at the Centro Nacional de Investigaciones Cardiovasculares (Madrid, Spain) in accordance with national and European Legislation. Procedures were approved by the CNIC Animal Welfare Ethics Committee and by the Area of Animal Protection of the Regional Government of Madrid (ref. PROEX 196/14).

## Author Contributions

Conceptualization: JV, MM; Methodology: JV, IR; Software: RR; Validation: JV, IR, RR, JA; Formal Analysis: JV, RR; Investigation: JV, IR, RR, JA; Data Curation: JV; Writing – Original Draft: JV, MM; Writing – Review & Editing: JV, IR, RR, JA, MM; Visualization: JV, MM; Supervision: JV, MM; Project Administration: JV, MM; Funding Acquisition: MM.

## Acknowledgements

We thank Maria Tiana for comments on the manuscript, and her and other members of the Manzanares lab for continued help and support, Diego Balboa and Timo Otonkoski for the gift of plasmids, Juan Bernal for plasmids and advice on using the piggyBac system, and Lindsey Montefiori and Marcelo Nobrega for help in analyzing their promoter-capture Hi-C data from human iPSC and iPSC-derived cardiomyocytes.

## Funding

This work was supported by the Spanish Ministerio de Ciencia e Innovación (grant BFU2017-84914-P to MM). JV was supported by a fellowship from the Fundación Bancaria “La Caixa” (LCF/BQ/DE15/10360011). The CBMSO is supported by an Institutional grant from the Fundación Ramon Areces, and the CNIC by the Instituto de Salud Carlos III (ISCIII), the Ministerio de Ciencia e Innovación and the Pro CNIC Foundation.

## Conflict of interest

The authors declare no conflict of interest

## Notes

### Competing Interest Statement

The authors have declared no competing interest.

## References

1. Schoenfelder, S. and Fraser, P. (2019) Long-range enhancer–promoter contacts in gene expression control. Nat. Rev. Genet., 10.1038/s41576-019-0128-0.

2. Gasperini, M., Tome, J.M. and Shendure, J. (2020) Towards a comprehensive catalogue of validated and target-linked human enhancers. Nat. Rev. Genet., 21, 292–310.

3. Visel, A., Minovitsky, S., Dubchak, I. and Pennacchio, L.A. (2007) VISTA Enhancer Browser - A database of tissue-specific human enhancers. Nucleic Acids Res., 10.1093/nar/gkl822.

4. Kvon, E.Z. (2015) Using transgenic reporter assays to functionally characterize enhancers in animals. Genomics, 10.1016/j.ygeno.2015.06.007.

5. Manzanares, M., Wada, H., Itasaki, N., Trainor, P.A., Krumlauf, R. and Holland, P.W.H. (2000) Conservation and elaboration of Hox gone regulation during evolution of the vertebrate head. Nature, 10.1038/35048570.

6. Banerji, J., Olson, L. and Schaffner, W. (1983) A lymphocyte-specific cellular enhancer is located downstream of the joining region in immunoglobulin heavy chain genes. Cell, 10.1016/0092-8674(83)90015-6.

7. Gillies, S.D., Morrison, S.L., Oi, V.T. and Tonegawa, S. (1983) A tissue-specific transcription enhancer element is located in the major intron of a rearranged immunoglobulin heavy chain gene. Cell, 10.1016/0092-8674(83)90014-4.

8. Mercola, M., Wang, X.F., Olsen, J. and Calame, K. (1983) Transcriptional enhancer elements in the mouse immunoglobulin heavy chain locus. Science (80-.).,221, 663–665.

9. Dunham, I., Kundaje, A., Aldred, S.F., Collins, P.J., Davis, C.A., Doyle, F., Epstein, C.B., Frietze, S., Harrow, J., Kaul, R., et al. (2012) An integrated encyclopedia of DNA elements in the human genome. Nature, 10.1038/nature11247.

10. Manolio, T.A., Collins, F.S., Cox, N.J., Goldstein, D.B., Hindorff, L.A., Hunter, D.J., McCarthy, M.I., Ramos, E.M., Cardon, L.R., Chakravarti, A., et al. (2009) Finding the missing heritability of complex diseases. Nature, 10.1038/nature08494.

11. Manolio, T.A. (2010) Genomewide Association Studies and Assessment of the Risk of Disease. N. Engl. J. Med.,363, 166–176.

12. Rickels, R. and Shilatifard, A. (2018) Enhancer Logic and Mechanics in Development and Disease. Trends Cell Biol., 10.1016/j.tcb.2018.04.003.

13. Tsai, C.T., Hsieh, C.S., Chang, S.N., Chuang, E.Y., Ueng, K.C., Tsai, C.F., Lin, T.H., Wu, C.K., Lee, J.K., Lin, L.Y., et al. (2016) Genome-wide screening identifies a KCNIP1 copy number variant as a genetic predictor for atrial fibrillation. Nat. Commun., 7, 10190.

14. Mullaney, J.M., Mills, R.E., Stephen Pittard, W. and Devine, S.E. (2010) Small insertions and deletions (INDELs) in human genomes. Hum. Mol. Genet., 10.1093/hmg/ddq400.

15. Smemo, S., Tena, J.J., Kim, K.H., Gamazon, E.R., Sakabe, N.J., Gómez-Marín, C., Aneas, I., Credidio, F.L., Sobreira, D.R., Wasserman, N.F., et al. (2014) Obesity-associated variants within FTO form long-range functional connections with IRX3. Nature, 10.1038/nature13138.

16. Gupta, R.M., Hadaya, J., Trehan, A., Zekavat, S.M., Roselli, C., Klarin, D., Emdin, C.A., Hilvering, C.R.E., Bianchi, V., Mueller, C., et al. (2017) A Genetic Variant Associated with Five Vascular Diseases Is a Distal Regulator of Endothelin-1 Gene Expression. Cell, 170, 522–533.e15.

17. Chugh, S.S., Havmoeller, R., Narayanan, K., Singh, D., Rienstra, M., Benjamin, E.J., Gillum, R.F., Kim, Y.H., McAnulty, J.H., Zheng, Z.J., et al.(2014) Worldwide epidemiology of atrial fibrillation: A global burden of disease 2010 study. Circulation, 129, 837–847.

18. Lubitz, S.A., Yin, X., Lin, H.J., Kolek, M., Smith, J.G., Trompet, S., Rienstra, M., Rost, N.S., Teixeira, P.L., Almgren, P., et al. (2017) Genetic risk prediction of atrial fibrillation. Circulation, 135, 1311–1320.

19. Bapat, A., Anderson, C.D., Ellinor, P.T. and Lubitz, S.A. (2018) Genomic basis of atrial fibrillation. Heart, 104, 201–206.

20. Gudbjartsson, D.F., Arnar, D.O., Helgadottir, A., Gretarsdottir, S., Holm, H., Sigurdsson, A., Jonasdottir, A., Baker, A., Thorleifsson, G., Kristjansson, K., et al. (2007) Variants conferring risk of atrial fibrillation on chromosome 4q25. Nature, 448, 353–357.

21. Benjamin, E.J., Rice, K.M., Arking, D.E., Pfeufer, A., Van Noord, C., Smith, A. V., Schnabel, R.B., Bis, J.C., Boerwinkle, E., Sinner, M.F., et al. (2009) Variants in ZFHX3 are associated with a trial fibrillation in individuals of European ancestry. Nat. Genet., 41, 879–881.

22. Ellinor, P.T., Lunetta, K.L., Glazer, N.L., Pfeufer, A., Alonso, A., Chung, M.K., Sinner, M.F., De Bakker, P.I.W., Mueller, M., Lubitz, S.A., et al. (2010) Common variants in KCNN3 are associated with lone atrial fibrillation. Nat. Genet., 42, 240–244.

23. Ellinor, P.T., Lunetta, K.L., Albert, C.M., Glazer, N.L., Ritchie, M.D., Smith, A. V., Arking, D.E., Müller-Nurasyid, M., Krijthe, B.P., Lubitz, S.A., et al. (2012) Meta-analysis identifies six new susceptibility loci for atrial fibrillation. Nat. Genet., 44, 670–675.

24. Low, S.K., Takahashi, A., Ebana, Y., Ozaki, K., Christophersen, I.E., Ellinor, P.T., Ogishima, S., Yamamoto, M., Satoh, M., Sasaki, M., et al. (2017) Identification of six new genetic loci associated with atrial fibrillation in the Japanese population. Nat. Genet., 49, 953–958.

25. Christophersen, I.E., Rienstra, M., Roselli, C., Yin, X., Geelhoed, B., Barnard, J., Lin, H., Arking, D.E., Smith, A. V., Albert, C.M., et al. (2017) Large-scale analyses of common and rare variants identify 12 new loci associated with atrial fibrillation. Nat. Genet., 49, 946–952.

26. Roselli, C., Chaffin, M.D., Weng, L.-C., Aeschbacher, S., Ahlberg, G., Albert, C.M., Almgren, P., Alonso, A., Anderson, C.D., Aragam, K.G., et al. (2018) Multi-ethnic genome-wide association study for atrial fibrillation. Nat. Genet., 10.1038/s41588-018-0133-9.

27. Nielsen, J.B., Thorolfsdottir, R.B., Fritsche, L.G., Zhou, W., Skov, M.W., Graham, S.E., Herron, T.J., McCarthy, S., Schmidt, E.M., Sveinbjornsson, G., et al. (2018) Biobank-driven genomic discovery yields new insight into atrial fibrillation biology. Nat. Genet.,50, 1234–1239.

28. Tam, V., Patel, N., Turcotte, M., Bossé, Y., Paré, G. and Meyre, D. (2019) Benefits and limitations of genome-wide association studies. Nat. Rev. Genet., 10.1038/s41576-019-0127-1.

29. Daly, M.J., Rioux, J.D., Schaffner, S.F., Hudson, T.J. and Lander, E.S. (2001) High-resolution haplotype structure in the human genome. Nat. Genet., 10.1038/ng1001-229.

30. Reich, D.E., Cargili, M., Boik, S., Ireland, J., Sabeti, P.C., Richter, D.J., Lavery, T., Kouyoumjian, R., Farhadian, S.F., Ward, R., et al. (2001) Linkage disequilibrium in the human genome. Nature, 10.1038/35075590.

31. Wall, J.D. and Pritchard, J.K. (2003) Haplotype blocks and linkage disequilibrium in the human genome. Nat. Rev. Genet., 10.1038/nrg1123.

32. Anderson, E.C. and Novembre, J. (2003) Finding haplotype block boundaries by using the minimum-description-length principle. Am. J. Hum. Genet., 10.1086/377106.

33. Cadiñanos, J. and Bradley, A. (2007) Generation of an inducible and optimized piggyBac transposon systemy. Nucleic Acids Res., 10.1093/nar/gkm446.

34. Aguirre, L.A., Alonso, M.E., Badía-Careaga, C., Rollán, I., Arias, C., Fernández-Miñán, A., López-Jiménez, E., Aránega, A., Gómez-Skarmeta, J.L., Franco, D., et al. (2015) Long-range regulatory interactions at the 4q25 atrial fibrillation risk locus involve PITX2c and ENPEP. BMC Biol., 13, 26.

35. Weltner, J., Balboa, D., Katayama, S., Bespalov, M., Krjutškov, K., Jouhilahti, E.M., Trokovic, R., Kere, J. and Otonkoski, T. (2018) Human pluripotent reprogramming with CRISPR activators. Nat. Commun., 10.1038/s41467-018-05067-x.

36. Gibson, D.G., Young, L., Chuang, R.Y., Venter, J.C., Hutchison, C.A. and Smith, H.O. (2009) Enzymatic assembly of DNA molecules up to several hundred kilobases. Nat. Methods, 10.1038/nmeth.1318.

37. Ran, F.A., Hsu, P.D., Wright, J., Agarwala, V., Scott, D.A. and Zhang, F. (2013) Genome engineering using the CRISPR-Cas9 system. Nat. Protoc., 10.1038/nprot.2013.143.

38. Concordet, J.P. and Haeussler, M. (2018) CRISPOR: Intuitive guide selection for CRISPR/Cas9 genome editing experiments and screens. Nucleic Acids Res., 10.1093/nar/gky354.

39. Yusa, K., Zhou, L., Li, M.A., Bradley, A. and Craig, N.L. (2011) A hyperactive piggyBac transposase for mammalian applications. Proc. Natl. Acad. Sci. U. S. A., 10.1073/pnas.1008322108.

40. Behringer, R., Gertsenstein, M., Vintersen Nagy, K. and Nagy, A. (2014) Manipulating the Mouse Embryo: A Laboratory Manual, Fourth Edition. Cold Harb. Lab. Press.

41. Van De Werken, H.J.G., Landan, G., Holwerda, S.J.B., Hoichman, M., Klous, P., Chachik, R., Splinter, E., Valdes-Quezada, C., Öz, Y., Bouwman, B.A.M., et al. (2012) Robust 4C-seq data analysis to screen for regulatory DNA interactions. Nat. Methods, 10.1038/nmeth.2173.

42. Gomez-Velazquez, M., Badia-Careaga, C., Lechuga-Vieco, A.V., Nieto-Arellano, R., Tena, J.J., Rollan, I., Alvarez, A., Torroja, C., Caceres, E.F., Roy, A., et al. (2017) CTCF counter-regulates cardiomyocyte development and maturation programs in the embryonic heart. PLoS Genet., 10.1371/journal.pgen.1006985.

43. Klein, F.A., Pakozdi, T., Anders, S., Ghavi-Helm, Y., Furlong, E.E.M. and Huber, W. (2015) FourCSeq: Analysis of 4C sequencing data. Bioinformatics, 10.1093/bioinformatics/btv335.

44. GTEx Consortium (2020) The GTEx Consortium atlas of genetic regulatory effects across human tissues. Science (80-.)., 10.1126/science.aaz1776.

45. Encode Project Consortium (2012) An integrated encyclopedia of DNA elements in the human genome. Nature.

46. Roadmap Epigenomics Consortium, Kundaje, A., Meuleman, W., Ernst, J., Bilenky, M., Yen, A., Heravi-Moussavi, A., Kheradpour, P., Zhang, Z., Wang, J., et al. (2015) Integrative analysis of 111 reference human epigenomes. Nature, 518, 317–329.

47. Ang, Y.-S., Rivas, R.N., Ribeiro, A.J.S., Srivas, R., Rivera, J., Stone, N.R., Pratt, K., Mohamed, T.M.A., Fu, J.-D., Spencer, C.I., et al. (2016) Disease Model of GATA4 Mutation Reveals Transcription Factor Cooperativity in Human Cardiogenesis. Cell, 167, 1734–1749.e22.

48. Luna-Zurita, L., Stirnimann, C.U., Glatt, S., Kaynak, B.L., Thomas, S., Baudin, F., Samee, M.A.H., He, D., Small, E.M., Mileikovsky, M., et al. (2016) Complex Interdependence Regulates Heterotypic Transcription Factor Distribution and Coordinates Cardiogenesis. Cell, 10.1016/j.cell.2016.01.004.

49. Kuhn, R.M., Haussler, D. and James Kent, W. (2013) The UCSC genome browser and associated tools. Brief. Bioinform., 10.1093/bib/bbs038.

50. Montefiori, L., Sobreira, D.R., Sakabe, N.J., Aneas, I., Joslin, A.C., Hansen, G.T., Bozek, G., Moskowitz, I.P., McNally, E.M. and Nóbrega, M.A. (2018) A promoter interaction map for cardiovascular disease genetics. Elife, 7.

51. Pennacchio, L.A., Ahituv, N., Moses, A.M., Prabhakar, S., Nobrega, M.A., Shoukry, M., Minovitsky, S., Dubchak, I., Holt, A., Lewis, K.D., et al. (2006) In vivo enhancer analysis of human conserved non-coding sequences. Nature, 444, 499–502.

52. Suzuki, S., Tsukiyama, T., Kaneko, T., Imai, H. and Minami, N. (2015) A hyperactive piggyBac transposon system is an easy-to-implement method for introducing foreign genes into mouse preimplantation embryos. J. Reprod. Dev., 10.1262/jrd.2014-157.

53. Lettice, L.A., Heaney, S.J.H., Purdie, L.A., Li, L., de Beer, P., Oostra, B.A., Goode, D., Elgar, G., Hill, R.E. and de Graaff, E. (2003) A long-range Shh enhancer regulates expression in the developing limb and fin and is associated with preaxial polydactyly. Hum. Mol. Genet., 10.1093/hmg/ddg180.

54. Furniss, D., Lettice, L.A., Taylor, I.B., Critchley, P.S., Giele, H., Hill, R.E. and Wilkie, A.O.M. (2008) A variant in the sonic hedgehog regulatory sequence (ZRS) is associated with triphalangeal thumb and deregulates expression in the developing limb. Hum. Mol. Genet., 10.1093/hmg/ddn141.

55. Shiratori, H., Sakuma, R., Watanabe, M., Hashiguchi, H., Mochida, K., Sakai, Y., Nishino, J., Saijoh, Y., Whitman, M. and Hamada, H. (2001) Two-step regulation of left-right asymmetric expression of Pitx2: Initiation by nodal signaling and maintenance by Nkx2. Mol. Cell, 10.1016/S1097-2765(01)00162-9.

56. Dixon, J.R., Selvaraj, S., Yue, F., Kim, A., Li, Y., Shen, Y., Hu, M., Liu, J.S. and Ren, B. (2012) Topological domains in mammalian genomes identified by analysis of chromatin interactions. Nature, 10.1038/nature11082.

57. Rada-Iglesias, A., Bajpai, R., Swigut, T., Brugmann, S.A., Flynn, R.A. and Wysocka, J. (2011) A unique chromatin signature uncovers early developmental enhancers in humans. Nature, 10.1038/nature09692.

58. Creyghton, M.P., Cheng, A.W., Welstead, G.G., Kooistra, T., Carey, B.W., Steine, E.J., Hanna, J., Lodato, M.A., Frampton, G.M., Sharp, P.A., et al. (2010) Histone H3K27ac separates active from poised enhancers and predicts developmental state. Proc. Natl. Acad. Sci. U. S. A., 10.1073/pnas.1016071107.

59. Claycomb, W.C., Lanson, N.A., Stallworth, B.S., Egeland, D.B., Delcarpio, J.B., Bahinski, A. and Izzo, N.J. (1998) HL-1 cells: A cardiac muscle cell line that contracts and retains phenotypic characteristics of the adult cardiomyocyte. Proc. Natl. Acad. Sci. U. S. A., 10.1073/pnas.95.6.2979.

60. Nadadur, R.D., Broman, M.T., Boukens, B., Mazurek, S.R., Yang, X., Van Den Boogaard, M., Bekeny, J., Gadek, M., Ward, T., Zhang, M., et al. (2016) Pitx2 modulates a Tbx5-dependent gene regulatory network to maintain atrial rhythm. Sci. Transl. Med., 8.

61. Lozano-Velasco, E., Hernández-Torres, F., Daimi, H., Serra, S.A., Herraiz, A., Hove-Madsen, L., Aránega, A. and Franco, D. (2016) Pitx2 impairs calcium handling in a dose-dependent manner by modulating Wnt signalling. Cardiovasc. Res., 109, 55–66.

62. Schlesinger, J., Schueler, M., Grunert, M., Fischer, J.J., Zhang, Q., Krueger, T., Lange, M., Tönjes, M., Dunkel, I. and Sperling, S.R. (2011) The cardiac transcription network modulated by gata4, mef2a, nkx2.5, srf, histone modifications, and microRNAs. PLoS Genet., 7.

63. Yeom, Y.I., Fuhrmann, G., Ovitt, C.E., Brehm, A., Ohbo, K., Gross, M., Hubner, K. and Scholer, H.R. (1996) Germline regulatory element of Oct-4 specific for the totipotent cycle of embryonal cells. Development, 122.

64. Tsai, C.T., Hsieh, C.S., Chang, S.N., Chuang, E.Y., Ueng, K.C., Tsai, C.F., Lin, T.H., Wu, C.K., Lee, J.K., Lin, L.Y., et al. (2016) Genome-wide screening identifies a KCNIP1 copy number variant as a genetic predictor for atrial fibrillation. Nat. Commun., 10.1038/ncomms10190.

65. Renou, L., Boelle, P.Y., Deswarte, C., Spicuglia, S., Benyoucef, A., Calvo, J., Uzan, B., Belhocine, M., Cieslak, A., Landman-Parker, J., et al. (2017) Homeobox protein TLX3 activates miR-125b expression to promote T-cell acute lymphoblastic leukemia. Blood Adv., 10.1182/bloodadvances.2017005538.

66. Nagpal, V., Rai, R., Place, A.T., Murphy, S.B., Verma, S.K., Ghosh, A.K. and Vaughan, D.E. (2016) MiR-125b Is Critical for Fibroblast-to-Myofibroblast Transition and Cardiac Fibrosis. Circulation, 10.1161/CIRCULATIONAHA.115.018174.

67. Mohenska, M., Tan, N.M., Tokolyi, A., Furtado, M.B., Costa, M.W., Perry, A.J., Hatwell-Humble, J., van Duijvenboden, K., Nim, H.T., Nilsson, S.K., et al. (2019) 3D-Cardiomics: A spatial transcriptional atlas of the mammalian heart. bioRxiv, 10.1101/792002.

68. Christophersen, I.E., Magnani, J.W., Yin, X., Barnard, J., Weng, L.C., Arking, D.E., Niemeijer, M.N., Lubitz, S.A., Avery, C.L., Duan, Q., et al. (2017) Fifteen Genetic Loci Associated with the Electrocardiographic P Wave. Circ. Cardiovasc. Genet., 10.1161/CIRCGENETICS.116.001667.

69. Parton, R.G. and Del Pozo, M.A. (2013) Caveolae as plasma membrane sensors, protectors and organizers. Nat. Rev. Mol. Cell Biol., 10.1038/nrm3512.

70. Lin, J., Lin, S., Choy, P.C., Shen, X., Deng, C., Kuang, S., Wu, J. and Xu, W. (2008) The regulation of the cardiac potassium channel (HERG) by caveolin-1. Biochem. Cell Biol., 10.1139/O08-118.

71. Langlois, S., Cowan, K.N., Shao, Q., Cowan, B.J. and Laird, D.W. (2008) Caveolin-1 and −2 interact with connexin43 and regulate gap junctional intercellular communication in keratinocytes. Mol. Biol. Cell, 10.1091/mbc.E07-06-0596.

72. Cheng, J.P.X., Mendoza-Topaz, C., Howard, G., Chadwick, J., Shvets, E., Cowburn, A.S., Dunmore, B.J., Crosby, A., Morrell, N.W. and Nichols, B.J. (2015) Caveolae protect endothelial cells from membrane rupture during increased cardiac output. J. Cell Biol., 10.1083/jcb.201504042.

73. Yang, K.C., Rutledge, C.A., Mao, M., Bakhshi, F.R., Xie, A., Liu, H., Bonini, M.G., Patel, H.H., Minshall, R.D. and Dudley, S.C. (2014) Caveolin-1 modulates cardiac gap junction homeostasis and arrhythmogenecity by regulating csrc tyrosine kinase. Circ. Arrhythmia Electrophysiol., 10.1161/CIRCEP.113.001394.

74. Williamson, I., Kane, L., Devenney, P.S., Flyamer, I.M., Anderson, E., Kilanowski, F., Hill, R.E., Bickmore, W.A. and Lettice, L.A. (2019) Developmentally regulated Shh expression is robust to TAD perturbations. Dev., 10.1242/dev.179523.

75. van Ouwerkerk, A.F., Bosada, F.M., van Duijvenboden, K., Hill, M.C., Montefiori, L.E., Scholman, K.T., Liu, J., de Vries, A.A.F., Boukens, B.J., Ellinor, P.T., et al. (2019) Identification of atrial fibrillation associated genes and functional non-coding variants. Nat. Commun., 10, 4755.

76. Smith, E. and Shilatifard, A. (2014) Enhancer biology and enhanceropathies. Nat. Struct. Mol. Biol., 10.1038/nsmb.2784.

77. Lupiáñez, D.G., Spielmann, M. and Mundlos, S. (2016) Breaking TADs: How Alterations of Chromatin Domains Result in Disease. Trends Genet., 10.1016/j.tig.2016.01.003.

78. Banerji, J., Rusconi, S. and Schaffner, W. (1981) Expression of a β-globin gene is enhanced by remote SV40 DNA sequences. Cell, 27, 299–308.

79. Moreau, P., Hen, R., Wasylyk, B., Everett, R., Gaub, M.P. and Chambon, P. (1981) The SV40 72 base repair repeat has a striking effect on gene expression both in SV40 and other chimeric recombinants. Nucleic Acids Res., 10.1093/nar/9.22.6047.

80. Nobrega, M.A., Ovcharenko, I., Afzal, V. and Rubin, E.M. (2003) Scanning Human Gene Deserts for Long-Range Enhancers. Science (80-.)., 10.1126/science.1088328.

81. Inoue, F. and Ahituv, N. (2015) Decoding enhancers using massively parallel reporter assays. Genomics, 10.1016/j.ygeno.2015.06.005.

82. Kvon, E.Z. (2015) Using transgenic reporter assays to functionally characterize enhancers in animals. Genomics, 106, 185–192.

83. Brinster, R.L., Chen, H.Y., Trumbauer, M.E., Yagle, M.K. and Palmiter, R.D. (1985) Factors affecting the efficiency of introducing foreign DNA into mice by microinjecting eggs. Proc. Natl. Acad. Sci. U. S. A., 10.1073/pnas.82.13.4438.

84. Brinster, R.L., Chen, H.Y., Trumbauer, M., Senear, A.W., Warren, R. and Palmiter, R.D. (1981) Somatic expression of herpes thymidine kinase in mice following injection of a fusion gene into eggs. Cell, 10.1016/0092-8674(81)90376-7.

85. Kawakami, K., Takeda, H., Kawakami, N., Kobayashi, M., Matsuda, N. and Mishina, M. (2004) A transposon-mediated gene trap approach identifies developmentally regulated genes in zebrafish. Dev. Cell, 10.1016/j.devcel.2004.06.005.

86. Bessa, J., Tavares, M.J., Santos, J., Kikuta, H., Laplante, M., Becker, T.S., Gómez-Skarmeta, J.L. and Casares, F. (2008) meis1 regulates cyclin D1 and c-myc expression, and controls the proliferation of the multipotent cells in the early developing zebrafish eye. Development, 10.1242/dev.011932.

87. Bessa, J., Tena, J.J., De La Calle-Mustienes, E., Fernández-Miñán, A., Naranjo, S., Fernández, A., Montoliu, L., Akalin, A., Lenhard, B., Casares, F., et al. (2009) Zebrafish Enhancer Detection (ZED) vector: A new tool to facilitate transgenesis and the functional analysis of cis-regulatory regions in zebrafish. Dev. Dyn., 10.1002/dvdy.22051.

88. Ivics, Z., Hackett, P.B., Plasterk, R.H. and Izsvák, Z. (1997) Molecular reconstruction of sleeping beauty, a Tc1-like transposon from fish, and its transposition in human cells. Cell, 10.1016/S0092-8674(00)80436-5.

89. Ruf, S., Symmons, O., Uslu, V.V., Dolle, D., Hot, C., Ettwiller, L. and Spitz, F. (2011) Large-scale analysis of the regulatory architecture of the mouse genome with a transposon-associated sensor. Nat. Genet., 10.1038/ng.790.

90. Uslu, V.V., Petretich, M., Ruf, S., Langenfeld, K., Fonseca, N.A., Marioni, J.C. and Spitz, F. (2014) Long-range enhancers regulating Myc expression are required for normal facial morphogenesis. Nat. Genet., 10.1038/ng.2971.

91. Symmons, O., Uslu, V.V., Tsujimura, T., Ruf, S., Nassari, S., Schwarzer, W., Ettwiller, L. and Spitz, F. (2014) Functional and topological characteristics of mammalian regulatory domains. Genome Res., 10.1101/gr.163519.113.

92. Symmons, O., Pan, L., Remeseiro, S., Aktas, T., Klein, F., Huber, W. and Spitz, F. (2016) The Shh Topological Domain Facilitates the Action of Remote Enhancers by Reducing the Effects of Genomic Distances. Dev. Cell, 10.1016/j.devcel.2016.10.015.

93. Shima, Y., Sugino, K., Hempel, C.M., Shima, M., Taneja, P., Bullis, J.B., Mehta, S., Lois, C. and Nelson, S.B. (2016) A mammalian enhancer trap resource for discovering and manipulating neuronal cell types. Elife, 10.7554/eLife.13503.

94. Kvon, E.Z., Zhu, Y., Kelman, G., Novak, C.S., Plajzer-Frick, I., Kato, M., Garvin, T.H., Pham, Q., Harrington, A.N., Hunter, R.D., et al. (2020) Comprehensive In Vivo Interrogation Reveals Phenotypic Impact of Human Enhancer Variants. Cell, 0, 1–10.

95. Ye, J., Tucker, N.R., Weng, L.C., Clauss, S., Lubitz, S.A. and Ellinor, P.T. (2016) A Functional Variant Associated with Atrial Fibrillation Regulates PITX2c Expression through TFAP2a. Am. J. Hum. Genet., 10.1016/j.ajhg.2016.10.001.

96. Zhang, M., Hill, M.C., Kadow, Z.A., Suh, J.H., Tucker, N.R., Hall, A.W., Tran, T.T., Swinton, P.S., Leach, J.P., Margulies, K.B., et al. (2019) Long-range Pitx2c enhancer–promoter interactions prevent predisposition to atrial fibrillation. Proc. Natl. Acad. Sci. U. S. A., 10.1073/pnas.1907418116.

97. Kirchhof, P., Kahr, P.C., Kaese, S., Piccini, I., Vokshi, I., Scheld, H.H., Rotering, H., Fortmueller, L., Laakmann, S., Verheule, S., et al. (2011) PITX2c is expressed in the adult left atrium, and reducing Pitx2c expression promotes atrial fibrillation inducibility and complex changes in gene expression. Circ. Cardiovasc. Genet., 10.1161/CIRCGENETICS.110.958058.

98. Chinchilla, A., Daimi, H., Lozano-Velasco, E., Dominguez, J.N., Caballero, R., Delpo, E., Tamargo, J., Cinca, J., Hove, L.M., Aranega, A.E., et al. (2011) PITX2 insufficiency leads to atrial electrical and structural remodeling linked to arrhythmogenesis. Circ. Cardiovasc. Genet., 10.1161/CIRCGENETICS.110.958116.

99. Reyat, J.S., Chua, W., Cardoso, V.R., Witten, A., Kastner, P.M., Nashitha Kabir, S., Sinner, M.F., Wesselink, R., Holmes, A.P., Pavlovic, D., et al. (2020) Reduced left atrial cardiomyocyte PITX2 and elevated circulating BMP10 predict atrial fibrillation after ablation. JCI Insight, 5.

100. Alvarez-Franco, A., Rouco, R., Ramirez, R.J., Guerrero-Serna, G., Tiana, M., Cogliati, S., Kaur, K., Saeed, M., Magni, R., Enriquez, J.A., et al. (2020) Transcriptome and proteome mapping in the sheep atria reveal molecular features of atrial fibrillation progression. Cardiovasc. Res., 10.1093/cvr/cvaa307.

101. Mizutani, S., Ishii, M., Hattori, A., Nomura, S., Numaguchi, Y., Tsujimoto, M., Kobayshi, H., Murohara, T. and Wright, J.W. (2008) New insights into the importance of aminopeptidase A in hypertension. Heart Fail. Rev., 10.1007/s10741-007-9065-7.

102. Gore-Panter, S.R., Hsu, J., Hanna, P., Gillinov, A.M., Pettersson, G., Newton, D.W., Moravec, C.S., Van Wagoner, D.R., Chung, M.K., Barnard, J., et al. (2014) Atrial Fibrillation Associated Chromosome 4q25 Variants Are Not Associated with PITX2c Expression in Human Adult Left Atrial Appendages. PLoS One, 9, e86245.

103. Sinner, M.F., Tucker, N.R., Lunetta, K.L., Ozaki, K., Smith, J.G., Trompet, S., Bis, J.C., Lin, H., Chung, M.K., Nielsen, J.B., et al. (2014) Integrating genetic, transcriptional, and functional analyses to identify 5 novel genes for atrial fibrillation. Circulation, 10.1161/CIRCULATIONAHA.114.009892.

104. Lee, J.Y., Kim, T.H., Yang, P.S., Lim, H.E., Choi, E.K., Shim, J., Shin, E., Uhm, J.S., Kim, J.S., Joung, B., et al. (2017) Korean atrial fibrillation network genome-wide association study for early-onset atrial fibrillation identifies novel susceptibility loci. Eur. Heart J., 10.1093/eurheartj/ehx213.

105. Thorolfsdottir, R.B., Sveinbjornsson, G., Sulem, P., Helgadottir, A., Gretarsdottir, S., Benonisdottir, S., Magnusdottir, A., Davidsson, O.B., Rajamani, S., Roden, D.M., et al. (2017) A Missense Variant in PLEC Increases Risk of Atrial Fibrillation. J. Am. Coll. Cardiol., 10.1016/j.jacc.2017.09.005.

106. Nielsen, J.B., Fritsche, L.G., Zhou, W., Teslovich, T.M., Holmen, O.L., Gustafsson, S., Gabrielsen, M.E., Schmidt, E.M., Beaumont, R., Wolford, B.N., et al. (2018) Genome-wide Study of Atrial Fibrillation Identifies Seven Risk Loci and Highlights Biological Pathways and Regulatory Elements Involved in Cardiac Development. Am. J. Hum. Genet., 10.1016/j.ajhg.2017.12.003.

107. Thorolfsdottir, R.B., Sveinbjornsson, G., Sulem, P., Nielsen, J.B., Jonsson, S., Halldorsson, G.H., Melsted, P., Ivarsdottir, E. V., Davidsson, O.B., Kristjansson, R.P., et al. (2018) Coding variants in RPL3L and MYZAP increase risk of atrial fibrillation. Commun. Biol., 10.1038/s42003-018-0068-9.

108. Benjamin, E.J., Rice, K.M., Arking, D.E., Pfeufer, A., Van Noord, C., Smith, A. V., Schnabel, R.B., Bis, J.C., Boerwinkle, E., Sinner, M.F., et al. (2009) Variants in ZFHX3 are associated with a trial fibrillation in individuals of European ancestry. Nat. Genet., 10.1038/ng.416.

109. Gudbjartsson, D.F., Holm, H., Gretarsdottir, S., Thorleifsson, G., Walters, G.B., Thorgeirsson, G., Gulcher, J., Mathiesen, E.B., Njølstad, I., Nyrnes, A., et al. (2009) A sequence variant in ZFHX3 on 16q22 associates with a trial fibrillation and ischemic stroke. Nat. Genet.,41, 876–878.

110. Ellinor, P.T., Lunetta, K.L., Glazer, N.L., Pfeufer, A., Alonso, A., Chung, M.K., Sinner, M.F., De Bakker, P.I.W., Mueller, M., Lubitz, S.A., et al. (2010) Common variants in KCNN3 are associated with lone atrial fibrillation. Nat. Genet., 10.1038/ng.537.

111. Lubitz, S.A., Lunetta, K.L., Lin, H., Arking, D.E., Trompet, S., Li, G., Krijthe, B.P., Chasman, D.I., Barnard, J., Kleber, M.E., et al. (2014) Novel genetic markers associate with atrial fibrillation risk in Europeans and Japanese. J. Am. Coll. Cardiol., 10.1016/j.jacc.2013.12.015.

112. Gudbjartsson, D.F., Helgason, H., Gudjonsson, S.A., Zink, F., Oddson, A., Gylfason, A., Besenbacher, S., Magnusson, G., Halldorsson, B. V., Hjartarson, E., et al. (2015) Large-scale whole-genome sequencing of the Icelandic population. Nat. Genet., 10.1038/ng.3247.

113. Pott, S. and Lieb, J.D. (2015) What are super-enhancers? Nat. Genet., 10.1038/ng.3167.

114. Arechederra, M., Carmona, R., González-Nuñez, M., Gutiérrez-Uzquiza, Á., Bragado, P., Cruz-González, I., Cano, E., Guerrero, C., Sánchez, A., López-Novoa, J.M., et al. (2013) Met signaling in cardiomyocytes is required for normal cardiac function in adult mice. Biochim. Biophys. Acta - Mol. Basis Dis., 10.1016/j.bbadis.2013.08.008.

115. Coutts, A.S., MacKenzie, E., Griffith, E. and Black, D.M. (2003) TES is a novel focal adhesion protein with a role in cell spreading. J. Cell Sci., 10.1242/jcs.00278.

116. Wang, Y., Botvinick, E.L., Zhao, Y., Berns, M.W., Usami, S., Tsien, R.Y. and Chien, S. (2005) Visualizing the mechanical activation of Src. Nature, 10.1038/nature03469.

117. Archacki, S.R., Angheloiu, G., Moravec, C.S., Liu, H., Topol, E.J. and Wang, Q.K. (2012) Comparative gene expression analysis between coronary arteries and internal mammary arteries identifies a role for the TES gene in endothelial cell functions relevant to coronary artery disease. Hum. Mol. Genet., 10.1093/hmg/ddr574.

118. Li, Y., Gao, S., Han, Y., Song, L., Kong, Y., Jiao, Y., Huang, S., Du, J. and Li, Y. (2020) Variants of Focal Adhesion Scaffold Genes Cause Thoracic Aortic Aneurysm. Circ. Res., 10.1161/CIRCRESAHA.120.317361.

119. Pang, B. and Snyder, M.P. (2020) Systematic identification of silencers in human cells. Nat. Genet., 52, 254–263.

120. Ngan, C.Y., Wong, C.H., Tjong, H., Wang, W., Goldfeder, R.L., Choi, C., He, H., Gong, L., Lin, J., Urban, B., et al. (2020) Chromatin interaction analyses elucidate the roles of PRC2-bound silencers in mouse development. Nat. Genet., 52, 264–272.

121. Berry, F.B., Miura, Y., Mihara, K., Kaspar, P., Sakata, N., Hashimoto-Tamaoki, T. and Tamaoki, T. (2001) Positive and Negative Regulation of Myogenic Differentiation of C2C12 Cells by Isoforms of the Multiple Homeodomain Zinc Finger Transcription Factor ATBF1. J. Biol. Chem., 10.1074/jbc.M010378200.

122. Jung, C.G., Kim, H.J., Kawaguchi, M., Khanna, K.K., Hida, H., Asai, K., Nishino, H. and Miura, Y. (2005) Homeotic factor ATBF1 induces the cell cycle arrest associated with neuronal differentiation. Development, 10.1242/dev.02098.

