## Supplementary Material for "Systematic in vivo interrogation identifies novel enhancers and silencers associated to Atrial Fibrillation"

**Victorino et al.**

**Supplementary Figures S1-S4**

**Supplementary Tables S1-S7**

**Table S1.** Cloning primers
**Table S2.** CRISPR guides
**Table S3.** RT-qPCR primers
**Table S4.** Primers for transcription of *piggyBac* mRNA
**Table S5.** Data of transgenic assays
**Table S6.** Primers used for genotyping of embryos
**Table S7.** 4C-seq primers

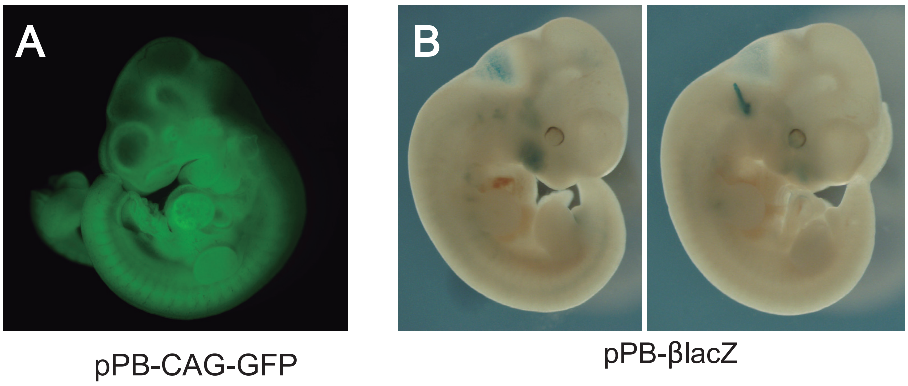


**Supplementary Figure S1. Efficient transgenesis in mouse embryos using the PB-ERA system. A**) Embryos microinjected with the pPB-CAG-GFP vector show high GFP signal with low degree of mosaicism. **B**) Examples of unspecific staining in E11.5 transgenic embryos carrying the empty vector (pPB-βlacZ).

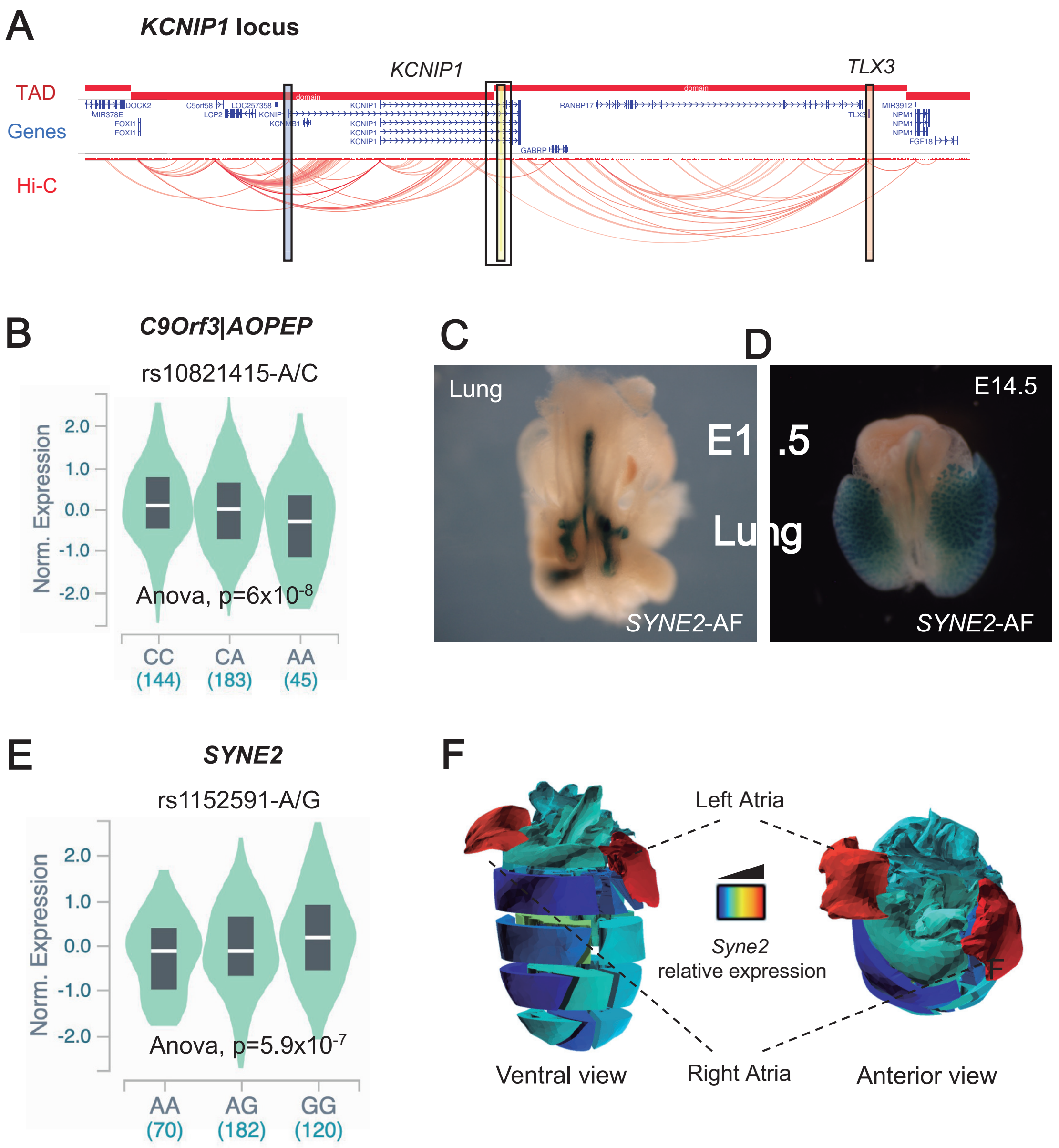


**Supplementary Figure S2. Candidate target genes of putative AF enhancers. A**) Schematic representation of the 1.5 Mb region (hg19; chr5:169,450,000-170,900,000) including *KCNIP1* and *TLX3*. Indicated on the left, from top to bottom, are topologically associated domains (TAD), annotated genes, and promoter-capture Hi-C data of hiPSC-derived cardiomyocytes (data from (50)). The *KCNIP1*-AF enhancer (yellow rectangle) is located at the boundary region between two consecutive TADs; nearby intronic regions (wider empty rectangle) interact with the promoters of *KCNIP1* (blue rectangle) and *TLX3* (orange rectangle). **B**) Transcriptomic data from GTEx (44) indicates that the risk allele (A) of rs10821415 correlates with lower expression of *C9orf3* in atrial tissue. **C-D**) Reporter expression driven by the *SYNE2*-AF element in lungs of E11.5 (**C**) and E14.5 (**D**) transgenic embryos. **E**) Transcriptomic data from GTEx (44) indicates that the risk allele (A) of rs1152591 correlates with lower expression of *SYNE2* in atrial tissue. **F**) Spatial representation of RNA-seq data from mouse heart tissue (3D-Cardiomics (67)) showing that *SYNE2* expression is highest in the atria.

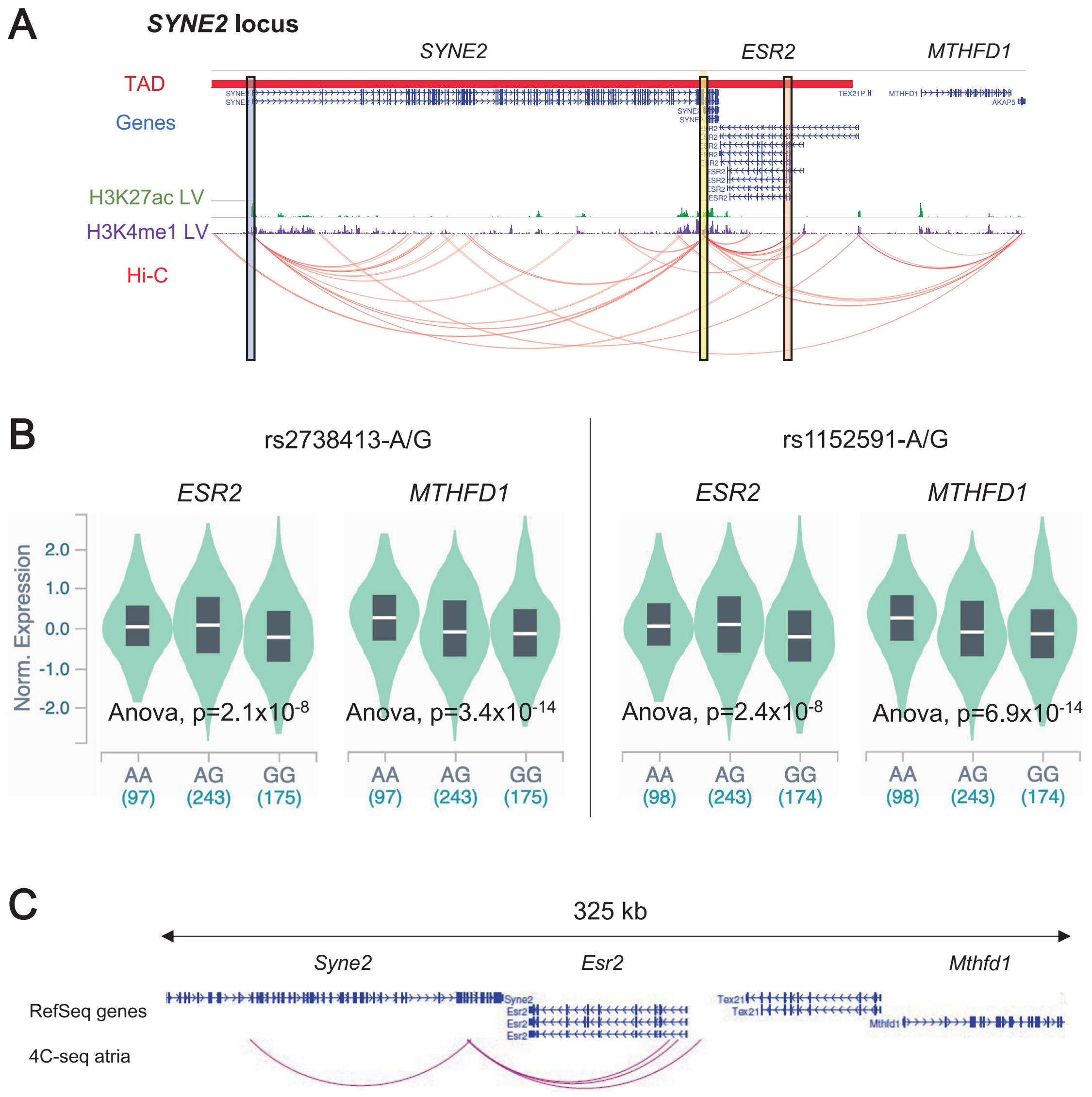


**Supplementary Figure S3. The SYNE2-AF enhancer regulates several genes. A**) Schematic representation of the 650 kb region (hg19; chr14:64,290,000-64,940,000) including *SYNE2*, *ESR2* and *MTHFD1*. Indicated on the left, from top to bottom, are topologically associated domains (TAD), annotated genes, ChIP-seq data for H3K27ac (green) and H3K4me1 (purple) in left ventricle (LV) from the Roadmap Epigenomics Project, and promoter-capture Hi-C data of hiPSC-derived cardiomyocytes (data from (50)). The *SYNE2*-AF region (yellow rectangle) interacts with the promoters of *SYNE2* (blue rectangle) and *ESR2* (orange rectangle). **B**) Risk alleles of AF variants rs2738413 and rs1152591 correlate with the expression of *ESR2* and *MTHFD1* genes in the lungs, where the enhancer is also active (Supplementary Figure S2C, D). **C**) 4C-seq in the atria of mouse hearts showing significant interactions in the 325 kb region (mm9; chr12:77,090,000-77,415,000) containing the orthologous region of *SYNE2*-AF.

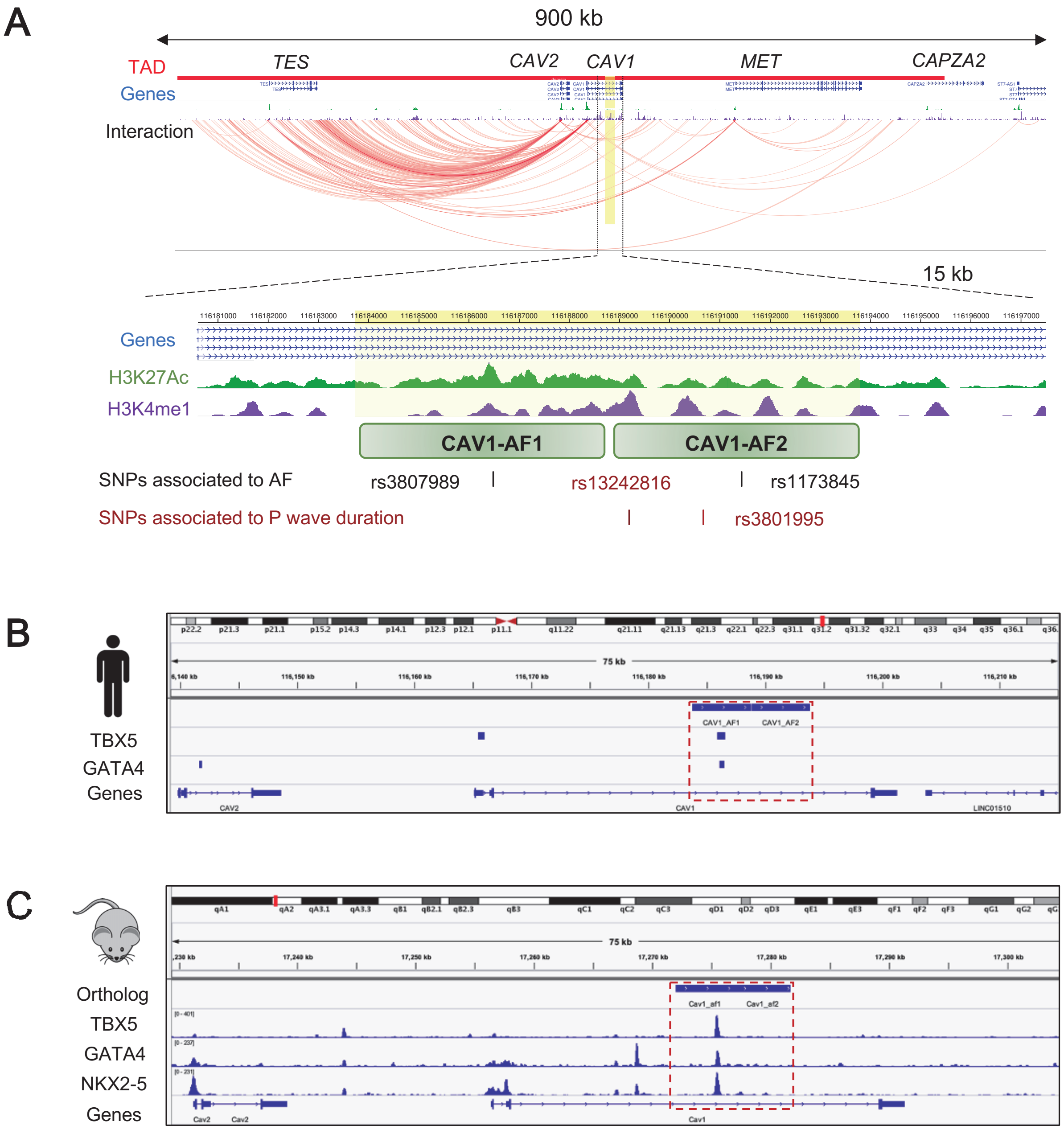


**Supplementary Figure S4. The 7q31 AF risk locus contains conserved epigenetic marks of regulatory elements. A**) Schematic representation of the 900 kb region (hg19; chr7:115,760,000-116,660,000) including *CAV1*, *CAV2*, *TES*, *MET* and *CAPZA2*. Indicated on the left, from top to bottom, are topologically associated domains (TAD), annotated genes, and promoter-capture Hi-C data of hiPSC-derived cardiomyocytes (data from (50)). The region containing *CAV1*-AF1 and *CAV1*-AF2 is indicated by a yellow rectangle. This 17 kb region (hg19; chr7:116,180,600-116,197,400) is shown below, including ChIP-seq data from the Roadmap Epigenomics Project for H3K27ac (green) and H3K4me1 (purple) in left ventricle (LV). Marks of active enhancers are enriched in the two candidate regions *CAV1*-AF1 and *CAV1*-AF2, where variants associated to AF (black) or P wave duration map. **B-C**) Genome browser views of 76 kb from the human (**B**; hg19; chr7:116,139,000-116,215,000) and the orthologous mouse region (**C**; mm9; chr6:17,229,500-17,305,000), where ChIP-seq data for cardiac TFs in hiPSC-derived cardiomyocytes (TBX5 and GATA4; data from (47)) and mESC (TBX5, GATA4 and NKX2-5; data from (48)) are shown. The red dashed rectangle indicates the enhancer region that is bound by the cardiac TFs in both organisms.
